# RNA 2’-OH modification with stable reagents enabled by nucleophilic catalysis

**DOI:** 10.1101/2025.09.05.674511

**Authors:** Eric T. Kool, Sumon Pratihar, Pavitra S. Thacker, Dipanwita Banerjee, Moon Jung Kim, Ryuta Shioi

## Abstract

RNA modification at 2’-OH has typically required highly reactive acylating species that exhibit short half-lives in water, challenging purification, and limited shelf lives. Here, we investigate the use of more stable species as electrophilic reagents, employing nucleophilic catalysis to promote reactions. Results show that multiple previously unreported electrophiles can react in high stoichiometric yields with RNA under appropriate catalysis. Most notably, aryl esters can transfer acyl groups to RNA in one hour, but are stable for months even in pure water. The results expand the functional chemotypes of RNA-reactive species, and identify reagent classes with improved stability and selectivity.

## Introduction

Posttranscriptional RNA modification at 2′-OH has undergone rapid growth in applications over the past decade, ranging from basic science applications such as probing secondary structure of folded RNAs,^1-5^ to practical methods for modifying messenger RNAs to improve potential therapeutic properties.^6^ Early studies (Fig. 1) documented the trace-level reactivity of isatoic anhydride and acylimidazole electrophiles with 2′-OH groups, enabling the development of widely useful methods for probing secondary structure, by taking advantage of the reactivity preference for unpaired nucleotides over those in double helical regions.^1-5^ While RNA structure probing typically requires only very low yields, many other applications require high yields for their utility.^7^ For example, conjugation of RNAs at 2′-OH with dyes and affinity reagents can be practically feasible with acylimidazole reagents, which can react in stoichiometrically high yields.^8^ Even higher levels of modification can enable useful applications in stabilising mRNAs, employing reactions and reagents where up to 90% of unpaired nucleotides are modified.^6^ Also contributing to the utility of these high-yield reactions is the development of strategies for site-localised RNA modification.^8^

**Fig. 1.**
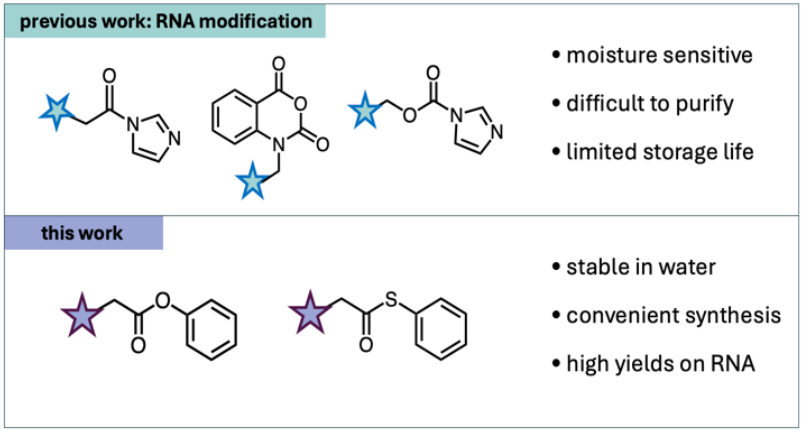
Reagents for high-yield RNA 2′-OH modification. Previous electrophilic reagents are moisture sensitive and can be difficult to purify and store. Reagents described here show high stability, with aqueous half-lives of years, while being highly reactive with RNA in the presence of a nucleophilic catalyst.

While this nonenzymatic approach for modifying RNA has found substantial utility, most of the reagents that are used for this purpose are highly reactive, exhibiting short half-lives in water and limited stability.^9^ For example, acylimidazole reagents can modify RNA in high yields at 2′-OH groups, but the reagents typically display half-lives in water of *ca*. 30-60 min, thus providing only a short window of reactivity. Yet more reactive are isatoic anhydride reagents, which can exhibit half-lives of seconds to minutes.^10^ This elevated reactivity can complicate the purification and storage of such reagents, as they typically cannot survive silica column chromatography, and their sensitivity to humidity limits their storage and use. Ideally, RNA-reactive groups would have extended stability against moisture while maintaining high reactivity for RNA.^11,12^ For these reasons, gaining a deeper understanding of the range of possible substrates for RNA reaction, and of catalysts that facilitate the reactions, may lead to more convenient and practical reagents, and improve researchers’ control over RNA modification for *in vitro* and cellular applications.

Here, we investigate the use of nucleophilic catalysis for high-yield modification of the 2′-OH groups of RNA using reduced-reactivity electrophiles that exhibit considerably greater stability to moisture than traditional acyl reagents. The chief aim is to develop more practical reagents for stoichiometric-level labelling and conjugation of RNA. Recent studies have observed that imidazole carbamate reagents can modify RNA under the influence of nucleophilic catalysis by N,N-dimethylaminopyridine (DMAP),^13-15^ which is a promising observation given the somewhat greater aqueous stability of such imidazole carbamate reagents relative to acylimidazoles (see below). A second class of electrophiles, anhydrides, can also be facilitated by DMAP in their RNA reactions,^16^ however those reagents are typically quite moisture-sensitive. Beyond this, the use of nucleophilic catalysis for RNA reaction has not been broadly surveyed with a wide range of acyl electrophiles. In addition, 2′-OH modification chemistries have recently expanded beyond acylation, to include sulphonylation and SNAr arylation as well; ^11,12,17^ while these reactions are mechanistically related to acylation, no studies of catalysis for these reactions with RNA have been reported.

Our experiments reveal that a broad range of electrophilic donors beyond imidazole carbamates can modify RNA in high yields with the aid of nucleophilic catalysis. A number of these reagents, such as aryl esters, were not known previously to react with RNA. In addition, we report that sulphonyl and aryl electrophiles can also modify RNA efficiently with nucleophilic catalysis. The results offer an expanded range of chemical versatility for high-yield conjugation in RNA research, and identify reactive species that are more stable for preparation, purification, and storage.

## Results

### Electrophile and catalyst survey

To examine the effects of nucleophilic catalysts on electrophilic modification of RNA at 2′-OH groups, we first performed a survey of acyl, sulphonyl, and aryl donors with a range of potential catalyst species^18^ (Fig. 2). For this initial survey, we included 13 electrophilic reagents and 6 potential catalysts along with uncatalysed controls (Fig. 2B,C), covering 91 combinations. Reactions for the survey were carried out in duplicate with an 18nt single-stranded RNA of mixed sequence in aqueous phosphate-saline buffer (pH 7.4) at room temperature for 6 h. The RNA contains 18 2′-OH groups that have the potential to react. In cases where previously unreported reactivity was observed, we tested whether the site of reaction was at 2′-OH groups as opposed to exocyclic amine groups of the nucleobases; this was done by performing control reactions using an 18mer DNA of identical sequence, which lacks 2′-OH groups but has the same exocyclic amines. Reactions were monitored by quantitative MALDI-TOF mass spectrometry, and we measured yields of RNA conversion (conversion of unreacted RNA to RNAs containing at least one modification) as well as median number of adducts on the RNA. A heat map summarising the survey is shown in Fig. 2C. The full data for each electrophile/nucleophile combination are given in the Supporting data file, along with representative MALDI-TOF spectra (Figs. S1-S3).

**Fig. 2.**
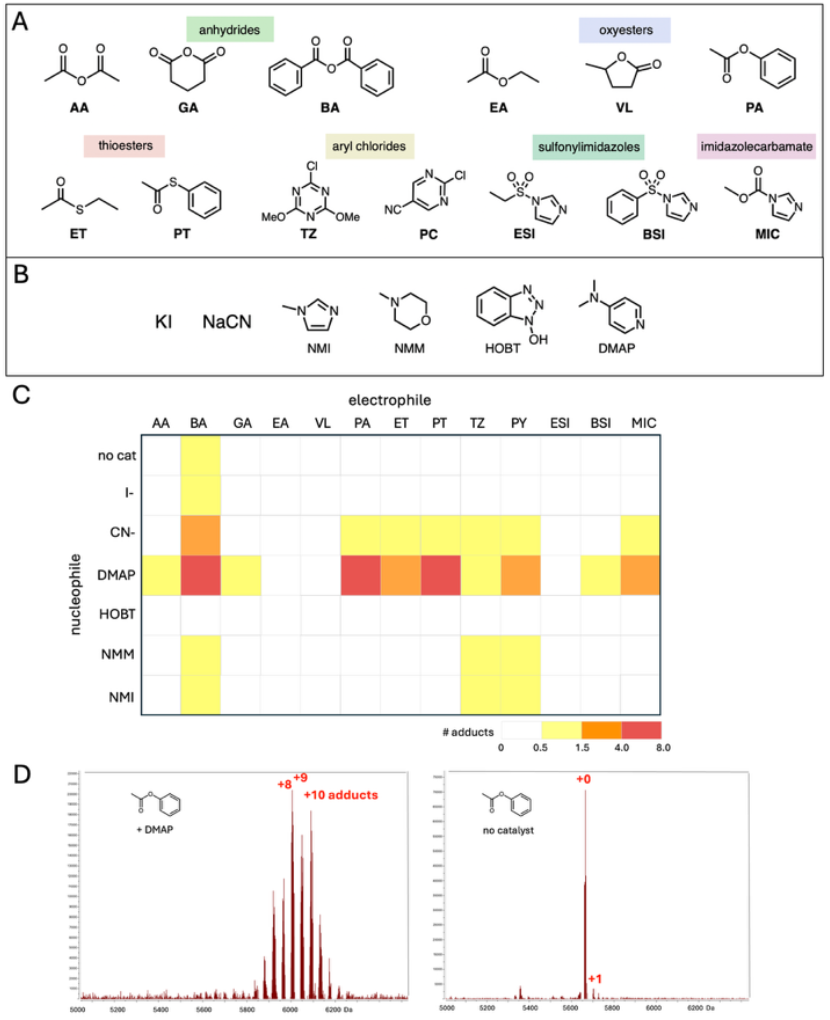
Reagents and nucleophiles tested in an initial survey. (A) Electrophilic reagents tested (abbreviations shown); (B) Nucleophiles tested; (C) Heat map of reactivity (as measured by RNA adduct numbers) for combinations of 13 reagents with 6 nucleophiles and buffer-only controls (“no cat”). Colour scheme indicates adduct numbers as shown. Reactions were performed in duplicate and results averaged. Reaction conditions: pH 7.4 phosphate buffered saline, 23°C, 6 h with 9 μM single-stranded RNA (18 nt), 40 mM reagent and 20 mM nucleophile. Adduct numbers were quantified by mass spectrometry (see SI for details); (D) Representative MALDI-TOF mass spectra of 18 nt single-stranded test RNA (MW=5650) reacted with phenyl acetate (PA) with and without DMAP (20 mM), showing an average of 8.5 acetyl adducts per RNA strand with the nucleophilic catalyst and a trace of acetylation (4% of a single adduct) without it.

The reactivity survey identified several successful electrophile and catalyst combinations that were previously unknown for RNA 2′-OH modification (Fig. 2C). Especially noteworthy were examples of oxyesters and thioesters, some of which were found to highly modify RNA with an effective catalyst. Under the reaction conditions with 20 mM DMAP, both alkyl and aryl thioacetates (**ET** and **PT**) transferred acetyl groups to RNA efficiently, adding multiple acyl groups to the test ssRNA. Perhaps more surprising was the observation that an aryl oxyester (phenyl acetate (**PA**)) reacted highly efficiently, adding as many as ten acetyl groups to the 18mer RNA strand in 6 h. The alkyl esters tested (ethyl acetate (**EA**) and γ-valerolactone (**VL**)) did not react to a significant extent. Also observed to react well was benzoic anhydride (**BA**), while the aliphatic anhydrides (acetic anhydride (**AA**) and glutaric anhydride (**GA**)) reacted moderately or only slightly, highlighting the additional reactivity conferred by the aromatic acyl group. **BA** also reacted to a significant extent with the RNA even in the absence of catalyst. Consistent with prior reports of imidazole carbamates reacting with RNA,^13-15,19,20^ we also found that methyl imidazole carbamate (**MIC**) transferred its acyl group moderately to RNA under DMAP catalysis, yielding an average of 1.8 carbonate adducts to each strand under these conditions.

Interestingly and somewhat surprisingly, previously untested classes of electrophiles also reacted with RNA with the assistance of nucleophilic catalysts. Two recent studies documented the reactivity of selected sulphonyl electrophiles with RNA 2′-OH groups.^12,17^ One report found that sulphonylimidazoles did not react well when tested alone,^12^ but here we found that the addition of DMAP promoted the reaction of benzenesulphonyl imidazole (**BSI**) with RNA, although the less electrophilic alkyl case (ethanesulphonyl imidazole, **ESI**) did not react measurably. Also noteworthy was the reactivity of SNAr reagents; we found that a pyrimidine aryl chloride (**PY**) and a triazene chloride (**TZ**) reacted poorly or not at all with RNA on their own as reported previously,^11^ but we found here that multiple nucleophilic catalysts were able to confer RNA reactivity to both these electrophiles. DMAP served as the most efficient catalyst, providing multiple aryl ether adducts on the test RNA.

While DMAP proved to be the most active nucleophilic catalyst in this survey, other species also provided enhancement of electrophilic reactions with RNA. Cyanide (20 mM) promoted the reaction of multiple electrophiles including an anhydride, aryl esters, an imidazole carbamate, and aryl chlorides (Figs. 2C and S1,2S). Tertiary amines (N-methylmorpholine and N-methylimidazole) also promoted reaction of multiple species, albeit less effectively than did DMAP.

Summarising the above data, we found that several classes of reagents, including multiple previously unreported functional groups, were responsive to RNA 2′-OH modification reactions in the presence of DMAP (Fig. 3A). The overall ranking of RNA reactivity for those catalyst-responsive functional groups was **PA** ∼ **PT** > **BA** > **MIC** > **BSI** ∼ **TZ**. Interestingly, comparisons of RNA modification yields in the presence and absence of DMAP shows (Fig. 3B) that the distinct classes of electrophiles exhibit marked differences in their catalyst responsiveness. Imidazole carbamates were previously used to modify RNA with DMAP,^13-15^ but our experiments with methyl imidazole carbamate (**MIC**) revealed only a modest 4-fold increase in adduct yields with catalyst (20 mM). An anhydride (**BA**) and an aryl chloride (**TZ**) showed more robust 14-to 21-fold increases in adduct yield with DMAP. Benzenesulphonylimidazole (**BSI**) also showed a considerable response, as its reactivity with RNA alone was quite low (estimated at 2% or less of a single adduct), but yields with DMAP were moderate. In contrast, the aryl esters phenyl thioacetate and phenyl acetate showed very large increases in reaction yields with the addition of DMAP, at 74- and 93-fold, respectively (Fig. 3B), and featured the highest yields of RNA adducts of the study. This finding was unexpected, as the aryl esters are inherently less electrophilic than several of the other species in the study.

**Fig. 3.**
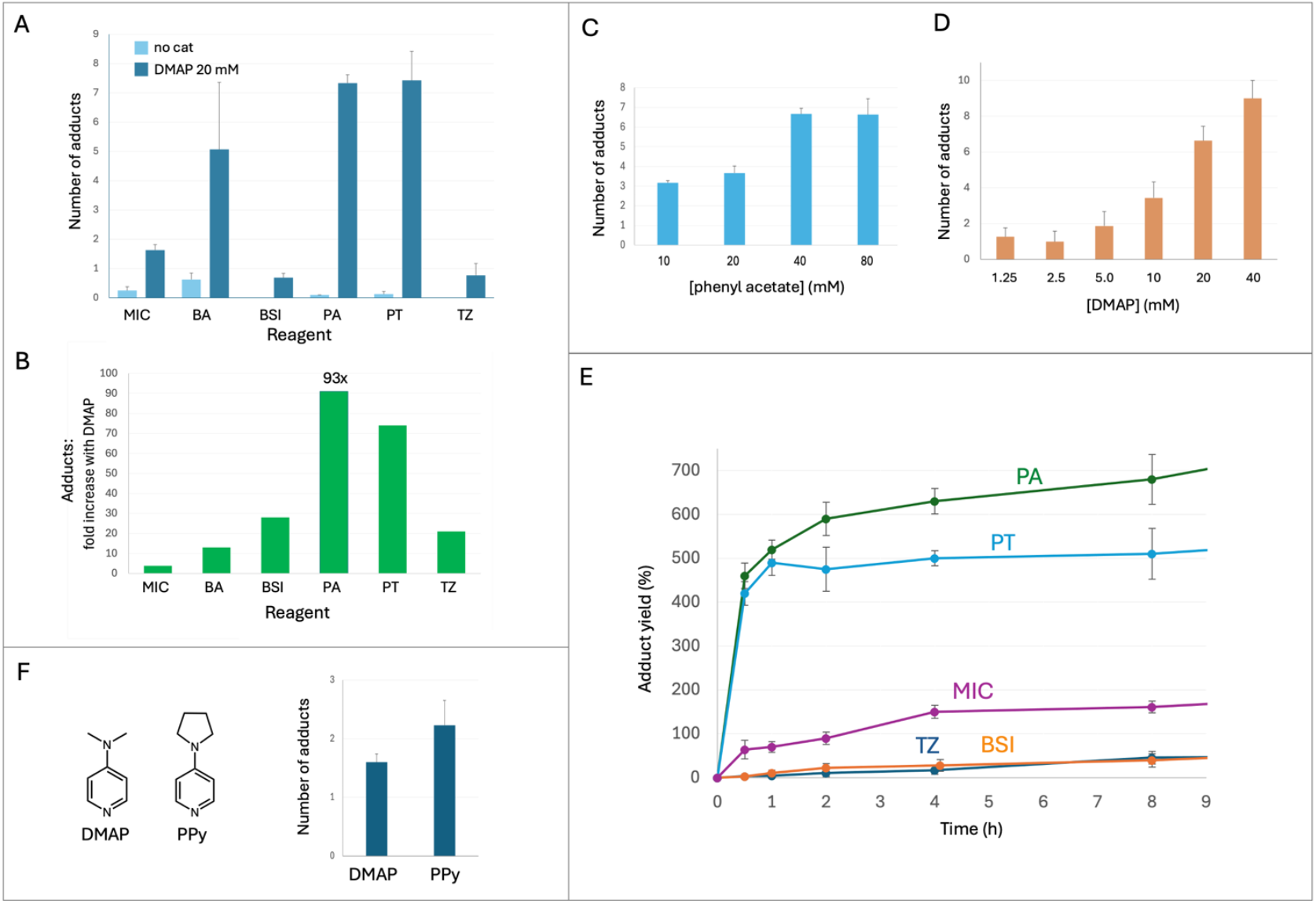
Effect of varied substrates and conditions on 4-aminopyridine-catalysed reaction with RNA. (A) Plot of adduct yields on a test 18 nt RNA strand for six classes of functional groups catalysed by DMAP; no-catalyst controls are also shown. See Fig. 2 for reagent abbreviations. Reaction conditions: 40 mM reagent, 20 mM catalyst, 9 mM RNA, 6 h, pH 7.4, 23 ^°^C. (B) Catalyst responsiveness for six reagents in (A), shown as fold increase in adducts per RNA strand with 20 mM DMAP added. (C) Effect of varied concentration of phenyl acetate on RNA ester yields ([DMAP]=20 mM); reactions were run for 2 h. (D) Effect of varied DMAP concentration ([**PA**]=40 mM) after 2 h; (E) Time courses of reaction with ssRNA for phenyl acetate **PA**, phenyl thioacetate **PT**, benzenesulphonylimidazole **BSI**, methyl imidazolecarbamate **MIC**, and chlorotriazine **TZ**. Yields are shown as percent RNA hydroxyl adducts. Conditions were as in (A); reactions were performed in quadruplicate with results averaged and error bars showing standard deviations. (E) Comparison of RNA yields for reaction with phenyl acetate catalysed by DMAP and by PPy. (A-D) Reactions were performed in triplicate with error bars showing standard deviations.

### Optimisation and scope of DMAP and analogue catalysis

Given the observation of broad catalysis of acyl transfer to RNA by DMAP, we explored the effect further with one of the most catalyst-responsive acyl agents observed here, phenyl acetate (**PA**). Ester reagent concentration varied from 10-80 mM revealed that 40 mM was sufficient for maximal yields (Figure 3C). Varying DMAP concentration resulted in increasing yields over the range 1.25-40 mM, with as low as 5 mM catalyst producing multiple adducts on the RNA after 2 h (Fig. 3D). We then compared time courses of reaction for five distinct classes of DMAP-responsive electrophiles that were identified with the above experiments. The data show (Fig. 3E) that phenyl acetate (**PA**) and phenyl thioacetate (**PT**) reacted considerably more rapidly than the other electrophiles, and reached higher yields, producing several acetyl adducts on the RNA after only 30 min. The reactions proceeded well beyond 100% RNA conversion by continuing to add more acetyl groups to the RNA, which contains 18 2′-OH groups in total. Although it is the most reactive case here, ultimately the reaction appears to plateau for reagent **PA**, reaching ester occupancy of about 2/3 of the 2′-OH positions. We interpret this tentatively as either the result of steric exclusion, or of dropping **PA** concentration as it is hydrolysed with the influence of DMAP. Interestingly, while reagent **PA** continued to add acetyl groups to the RNA in hours 1-12, the reaction with **PT** appeared to stall after one hour, possibly due to more rapid hydrolysis of the thioester reagent catalysed by DMAP. In contrast to these aryl ester reagents, **TZ, BSI** and **MIC** reacted considerably more slowly and provided limited yields even after 14 h (Fig. S4).

Next, we compared catalysts DMAP and an aminopyridine analogue reported to be more active in small-molecule reactions (4-pyrrolopyridine, PPy)^21^ for their relative ability to promote the reaction of phenyl acetate with RNA, and found that the latter conferred yet greater reactivity at 10 mM catalyst, promoting the addition of 2.2 acetyl groups on average to the RNA in 2 h, relative to 1.6 groups for the more traditional catalyst (Figs. 3F and S5). We then tested this more active catalyst with one of the least reactive substrates in the survey, ethyl acetate (**EA**). Remarkably, even this poorly reactive alkyl ester, typically considered an inert solvent, was able to acetylate RNA to a moderate extent in the presence of PPy (Fig. S6).

In the case of the ester acyl transfer reagents in this work, the products are ester adducts on the RNA. We wondered whether DMAP might also catalyse the hydrolysis of an ester adduct from the RNA over extended time, potentially affecting yields. To test this, we prepared multi-acetylated RNA using phenyl acetate. We then isolated the multi-acetylated RNA and incubated it with DMAP (10 mM) or buffer control over 24 h in the absence of phenyl acetate. The data show (Fig. S7) that only small losses of acetyl groups (*ca*. 25%) occurred after 24 h with 10 mM DMAP, as compared with the acylated RNA control. Thus we conclude that DMAP (if left in contact with the RNA product for extended time) may slowly catalyse the hydrolytic removal of acetyl esters from RNA, similar to results with other acyl adducts reported previously.^6^ However, we note that no acetyl loss is seen at up to 14 h when excess phenyl acetate is present (Fig. S4), as acetyl groups continue to be added.

### Stability of reagents

One goal of this study was to identify electrophilic RNA-modifying reagents with greater stability to moisture than prior isatoic anhydrides and acylimidazoles, which typically have half-lives of only minutes to an hour in water.^7^ Improving stability to water would make future RNA-modifying reagents simpler to purify, handle, and store than previous reagents. The above studies identified multiple new species that react well with RNA; to explore their relative hydrolytic stabilities, we measured half-lives of six reagents in D_2_O by NMR spectroscopy. Kinetics fits are given in Fig. S8, and half-lives are listed in Fig. 4A. The results show a wide range of aqueous stabilities. The shortest half-life (*ca*. 8 h) was seen for benzoic anhydride, while benzenesulphonyl imidazole and methyl imidazole carbamate were found to be more moderately sensitive to moisture, displaying greater stability (∼2-5 days). By comparison, the chloropyrimidine arylating agent is considerably more stable to hydrolysis (t_1/2_ ∼3 months), consistent with its distinct SNAr mode of reactivity. In contrast, phenyl thioacetate and phenyl acetate are far more stable than all other reagents measured here, showing only slight hydrolysis (4-6%) after one month at room temperature, with half-lives extrapolated to 1.5 and 2.7 years. Remarkably, the aryl ester reagents exhibit > 10^4^-fold improvement in stability to moisture over previously known RNA-modifying functional groups such as standard acylimidazole reagent NAI (t_1/2_ ∼ 30 min), ^4^ and >200-fold over methyl imidazolecarbamate. However, under DMAP catalysis, phenyl acetate reacts with RNA the most efficiently of the group (Figs. 3A,B).

**Fig. 4.**
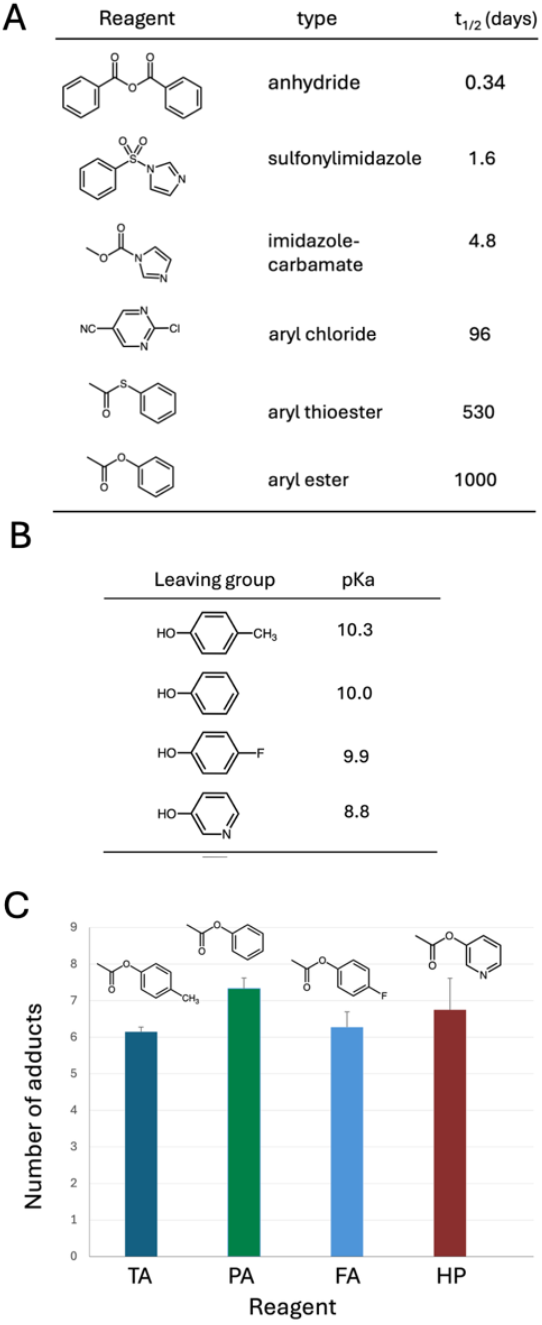
Stability and reactivity of DMAP-responsive RNA-modifying reagents. (A) Sensitivity to water (shown as half-lives) of six classes of reagents measured in D_2_O solution (+15 or 30% DMSO, see SI) at room temperature; (B) Varied phenolic leaving groups employed to test the effect of leaving group ability (pK_a_ values shown); (C) Graph of aryl ester reactivity with leaving groups of varied pK_a_. Graph shows number of RNA ester adducts at 6 h ([reagent]=40 mM) in the presence of 20 mM DMAP. Reactions were performed in triplicate with error bars showing standard deviations.

### Generality and selectivity of esters as substrates

The above data showed that aryl esters can act as especially efficient reaction substrates for RNA, and were not known previously to react with the biopolymer. To test whether small changes in leaving group ability can affect reactivity, we performed experiments to compare the reactivity of four aryl esters constructed from different phenolic groups having varied pK_a_ values ranging from 8.8-10.3 (Fig. 4B). Reactions in the presence of DMAP revealed that there was little difference in RNA reaction yields among the aryl ester variants, all producing a similarly robust number of adducts on the RNA (Fig. 4C).

### RNA modification with biological thioester Acetyl CoA

While aryl esters and thioesters were found to be highly reactive toward RNA under DMAP catalysis, our experiments also showed that an alkyl thioester (ethyl thioacetate) was a moderately reactive substrate as well, producing more RNA adducts than did an imidazole carbamate reagent (Fig. S2). Since thioesters are common electrophiles for biological acylation, we were prompted to ask whether the cofactor Acetyl CoA can modify RNA using DMAP as catalyst (Fig. S9). The results showed that multiple acetyl groups were transferred to the 18nt test RNA over 24 h.

### Testing phenyl esters of RNA-labelling agents

The above studies revealed that phenyl esters provided efficient RNA reactivity in the presence of millimolar DMAP, while being quite stable in the absence of catalyst. This suggests the possibility of more general utility of phenyl esters for RNA high-yield modifications. We explored this in a preliminary way by preparing phenyl esters of two common RNA labels, biotin and a diethylaminocoumarin dye. They were purchased in carboxylic acid form and coupled to phenol in a single step with EDC (Fig. 5; see ESI for details). We found that the labelling reagents could be readily purified by silica column chromatography, which is difficult or impossible for prior acylimidazole reagents.

**Fig. 5.**
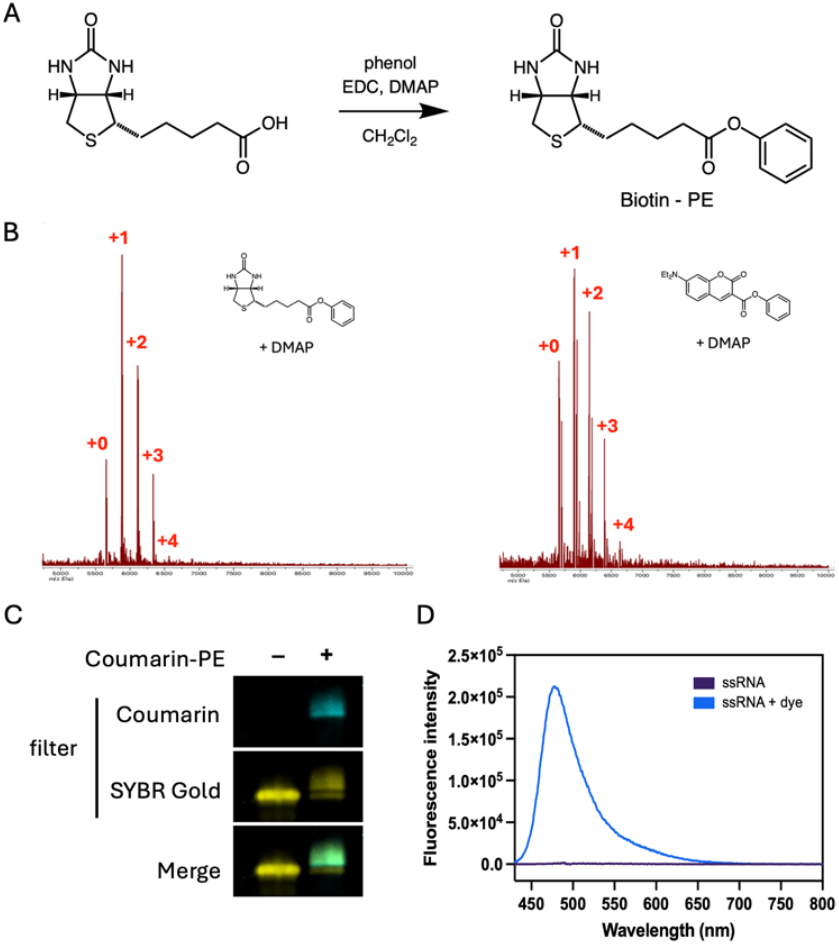
Labelling RNA with phenyl ester-derivatised labels biotin-PE and diethylaminocoumarin-PE. (A) One-step synthesis employed for preparation of reagents (biotin example is shown); (B) Mass spectra confirming RNA modified with biotin (left) and coumarin (right); (C) Fluorescence imaging of coumarin-labelled 18 nt RNA in polyacrylamide gel; (D) Fluorescence emission spectrum of coumarin-labelled RNA (λ_ex_ = 420 nm) isolated after reaction in comparison with untreated RNA; see ESI for details of synthesis and characterisation.

With these activated phenyl ester reagents in hand, we examined their ability to label a test ssRNA. Experiments confirmed that both reagents could label RNA effectively in the presence of DMAP, producing 79-85% yield of the RNA conjugates confirmed by mass spectrometry and, for the fluorophore, by imaging and spectroscopy of the modified RNA (Figs. 5C,D, S10,S11).

## Discussion and Conclusions

We have shown that a number of electrophilic reagent classes that are relatively stable to moisture can act as efficient acyl and aryl donors for RNA 2′-OH groups, by use of nucleophilic catalysis to promote reactivity. Multiple new categories of reactants, including aryl esters, thioesters, aryl sulphonylimidazoles, and aryl chlorides are all found to be responsive to nucleophilic catalysis in water with RNA as the acceptor. While imidazole carbamates were previously shown to react with RNA via DMAP catalysis,^13-16^ the new work establishes that several other reagent classes, some of them much more stable, can also benefit from this catalysis as well. We expect that these findings will result in multiple novel applications for RNA modification and labelling. High-yield reactions at 2′-OH have been applied to RNA caging, protection of mRNAs against degradation, intracellular delivery, RNA purification, and fluorescent labelling, and the new electrophilic groups may find utility in several of these areas.^9^ Although the current work demonstrates random-site modification of the RNA, such general reactivity at 2′-OH groups is useful both in caging of RNAs^13-16,19^ as well as in protection against RNA strand cleavage.^6^ In addition, we anticipate that the current reagents may also be applied to previously developed site-localised RNA modification methods as well.^8^

Most notably, our data document that phenolic esters are highly reactive with RNA with the aid of DMAP, and in fact are the most RNA-reactive of the electrophiles tested here in the presence of this nucleophilic catalyst. This is surprising, given the fact that a phenol ester is among the least electrophilic compounds in its reactivity to water without catalyst (Fig. 3A).

Mechanistic studies in organic solvent indicate that the rate-limiting step of DMAP transacylation catalysis is the attack of the alcohol nucleophile on the acylpyridinium intermediate.^22^ In this light, we observe considerably different rates of RNA reaction for three acetyl donors (ethyl acetate << acetic anhydride << phenyl acetate (Figs. S1,2)), all of which involve the same acetylpyridinium intermediate reacting with RNA. This suggests that the high RNA reactivity of phenyl acetate results from its ability to generate the acylpyridinium intermediate at higher concentrations than do the other two reagents. Further studies will be required to shed light on the special reactivity of the aryl esters in this case.

From the practical standpoint, our finding that aryl esters are efficient reactants for RNA is especially noteworthy, as they display far greater stability to moisture than all prior reagents that have been used to modify RNA.^9^ Indeed, with respect to prior RNA-reactive electrophile classes, we observed major loss of activity for benzoic anhydride and methyl imidazole carbamate stocks during these experiments (Fig. S12). In marked contrast, our data shows extended stability of the aryl ester reagents even in water solution, with half-lives of years. This establishes aryl esters especially as a generally stable class of RNA-modifying reagent that suffers far less moisture sensitivity than common RNA-modifying reagents such as acylimidazoles, imidazolecarbamates, and anhydrides, rendering them much more convenient to purify, handle, and store. The poor stability of the previous moisture-sensitive reagents can also be detrimental to experimental reproducibility, and may hinder application by non-specialist users.

Although we find that an aryl thioester, like the aryl oxyesters, is also highly reactive for RNA modification in the presence of DMAP, we anticipate that oxyesters of phenol may prove more practically useful for high-yield RNA modification given the oxidative instability and noxious odour of thiophenol, which is required for preparation of the former species. In any case, we show that phenyl esters of standard labelling agents are readily prepared in one step from commercial carboxylic acid derivatives, and can be purified by silica chromatography unlike more reactive species. One potential limitation of phenyl esters is their relative hydrophobicity; for example, biotin phenyl ester, having a substantial alkane chain (see Fig. 5A), displayed limited solubility in reactions at 40 mM (see SI). One alternative worthy of future exploration is the 3-hydroxypyridine leaving group, which confers similar reactivity as phenol (Fig. 4C) but may provide greater solubility.

The current work focuses on *in vitro* catalyst-promoted RNA modification rather than biological application, which will await the development of small-molecule catalysts that are biocompatible. However, our finding that the biological cofactor Acetyl-CoA can act as an acetyl donor to RNA 2′-OH groups in the presence of a catalyst suggests the possibility that this modification may occur in living cells assisted by enzymatic catalysis. Cellular RNAs are known to be acetylated at N4 of cytidine as a posttranscriptional modification,^23^ but the current results suggest that it is also biochemically plausible that 2′-OH modification has the potential to exist as well.

Our observation that, in addition to aryl esters, aryl chlorides and an aryl sulphonylimidazole can also undergo catalysis by DMAP and other nucleophiles with RNA as an acceptor has not been reported previously. Nucleophilic catalysis of SNAr reactivity is rare in the literature; we are aware of only one prior study that documented DMAP catalysis of SNAr reactivity of a heterocycle,^24^ and it required organic solvent, elevated temperature, and strong base to increase alcohol reactivity. We also know of no previous documentation of tertiary amine catalysis of SNAr reactions. We expect that this catalysis may prove useful in multiple schemes for site-localised RNA labelling.^8,25^ As for sulphonyl species, sulphonyl chlorides have been reacted with small molecule amines aided by DMAP in organic solvent,^26^ but we do not know of examples of this catalysis with the less-reactive sulphonylimidazoles. Although they are less stable than the above phenyl esters, both aryl chlorides and sulphonylimidazoles show considerably reduced sensitivity to moisture (see Fig. 4A) relative to earlier more highly reactive electrophiles for RNA such as anhydrides and acylimidazoles.

## Author contributions

E. T. K. conceived and directed the project and analysed data. E. T. K., S. P., P. T., D. B., M. J. K., and R. S. performed experiments. E. T. K. wrote the manuscript.

## Data availability

The data supporting this article have been included as part of the Supporting file.

## Acknowledgements

We thank the U.S. National Institutes of Health for support (GM145357).

## Notes and references

§ Equal contributions.

## Graphical abstract

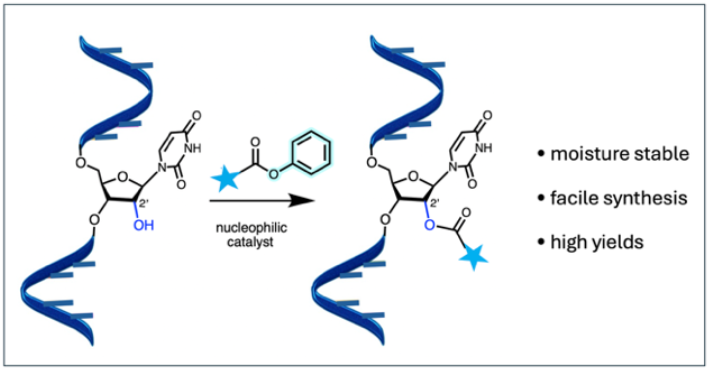

