## Supporting Information for "RNA 2’-OH modification with stable reagents enabled by nucleophilic catalysis"

### Materials

#### Table S1. Reagents and materials

| REAGENTS | SOURCE | IDENTIFIER |
|---|---|---|
| **Reagents and Enzymes** | | |
| 5M NaCl | Invitrogen | # AM9760G |
| 1M $MgCl_2$ | Invitrogen | # AM9530G |
| SequaGel - UreaGel Concentrate | National diagnostics | # EC-830 |
| SequaGel - UreaGel Diluent | National diagnostics | # EC-840 |
| SequaGel - UreaGel Buffer | National diagnostics | # EC-835 |
| Deuterium oxide | Sigma-Aldrich | # 151882 |
| Dimethyl sulfoxide-d6 | Sigma-Aldrich | # 151874 |
| Chloroform-d | Sigma-Aldrich | # 151823 |
| DMSO anhydrous | Sigma-Aldrich | # 276855 |
| Methanol-d4 | Sigma-Aldrich | # 422878 |
| gamma-butyrolactone | Millipore Sigma | # B103608-25 |
| Methyl imidazole carbamate | Enamine | EN300-7613220 |
| Benzoic anhydride | Fisher Scientific | # A14269.18 |
| Biotin | Millipore Sigma | #B4501-100 |
| 7-diethylaminocoumarin-3-carboxylic acid | Lumiprobe | none |
| p-tolyl acetate | Millipore Sigma | # A067025g |
| Phenyl nicotinate | Millipore Sigma | # S370355 |
| Phenyl acetate | Sigma-Aldrich | # 108723 |
| Phosphate, 0.5M buffer pH 7.4 | ThermoFisher | #J60785.AK |
| Phosphate Buffered Saline | Sigma-Aldrich | # P3813 |
| 2-Chloro-5-pyrimidinecarbonitrile | AA Blocks | # AA0020SZ |
| Glutaric anhydride | TCI America | # G0071 |
| Acetic anhydride, ACS reagent | Millipore Sigma | # 242845-5G |
| potassium iodide | Sigma-aldrich | #221945-5g |
| 1-Hydroxybenzotriazole Monohydrate | TCI America | #H0468 |
| 4-pyrrolidino-pyridine | Sigma-Aldrich | # 213373 |
| 2-Chloro-4,6-dimethoxy-1,3,5-triazine | Sigma-Aldrich | # 375217 |
| 4-Methylmorpholine | Sigma-Aldrich | # 407704 |
| Ethyl acetate | Sigma-Aldrich | # 319902 |
| S-phenyl thioacetate | ThermoFisher | # H55175.06 |
| 4-(Dimethylamino)-pyridine | Sigma-Aldrich | # 522821 |
| Methyl sulfoxide, 99.7+%, Extra Dry, AcroSeal™ | Thermo Scientific | # AC326881000 |
| Ammonium citrate dibasic | Sigma-Aldrich | # 09833 |
| 2′,4′,6′-Trihydroxyacetophenone monohydrate | Sigma-Aldrich | # T64602 |
| 1xPBS, pH=7.4 | gibco | # 10010-023 |
| UltraPure DNase/RNase-free Distilled water | Thermo Scientific | # 10977023 |
| Glycogen, RNA grade | Thermo Scientific | # R0551 |
| 3 M sodium acetate, pH=5.2 | Thermo Scientific | # AM9740 |
| 96% Ethanol | Fisher Scientific | # BP8202-500 |
| 70% Ethanol Solution | Fisher Scientific | #BP8201-1 |

**Table S2. Oligonucleotides used in this work** (purchased from IDT)

| Name | Sequence (5' -> 3') |
|---|---|
| **RNA oligonucleotide** | |
| Test 18 nt ssRNA | AUCCUGCCGACUACGCCA (MW: 5650 Da) |
| **DNA oligonucleotide** | |
| Test 18 nt ssDNA | ATCCTGCCGACTACGCCA (MW: 5404 Da) |

### Experimental Procedures

**RNA Reactions:** In a sterile 200 µL PCR tube, 5.0 µL of PBS buffer stock (3.3x), was mixed with 6.5 µL of 20.0 µM 18 nt test ssRNA aqueous stock solution. Electrophile reagents were prepared as stock solutions in anhydrous DMSO at 400 mM, and 1.5 µL of the stock solution of desired reagent was added to the reaction mixture, followed by 1.5 mL of catalyst stock solution (200 mM in 1:1 DMSO/nuclease-free water). These reactions were incubated for 2 or 6 h (variable for time courses) at 23°C, and the reacted RNA was subsequently isolated by ethanol precipitation. The level of RNA modification was measured by quantitative MALDI-TOF M/S analysis.
For selected reactions, parallel reactions were performed under identical conditions with 18 nt test ssDNA, a DNA oligonucleotide containing the same sequence as the RNA.

**Ethanol precipitation of RNA reactions:** For each RNA reaction, 14 µL reaction mixture was added to a precipitation solution composed of 117 µL RNAse-free water, 16.5 µL sodium acetate solution (0.33 M NaOAc (pH 5.2)) and 1.5 µL glycogen solution (20 mg/mL in water) and mixed by vortexing. 450 µL of ice-cold 96% ethanol was then added, and the mixture mixed by vortexing. After cooling at -80°C for at least 30 min, the mixture was centrifuged at 14.4k RPM for 14 min at 4°C. The supernatant was discarded to obtain a pellet, which was washed with 70% ice-cold ethanol twice. The obtained pellet was air dried for 30 min and subsequently dissolved in 2.5 µL water for direct measurement with mass spectrometry or in further experiments (below).

**Biotin-phenyl ester conjugation:** In a sterile 200 µL PCR tube, 6.5 µL 18nt test ssRNA (20 µM) was combined with 6.0 µL 3.3x PBS buffer containing 15 mM $MgCl_2$, 3.9 µL water, 1.5 µL DMAP stock (200 mM in 50% aqueous DMSO), and 1.5 µL Biotin Phenyl Ester (biotin-PE) stock in DMSO (400 mM). (Note: upon addition of the biotin stock, a white precipitate was observed). The mixture was incubated for 6 h at 23°C with occasional vortexing. The mixture was then subjected to ethanol precipitation to isolate the RNA and remove unreacted biotin-PE reagent. The obtained pellet was then dissolved in 2.5 µL nuclease-free water for mass spectrometric analysis.

**Coumarin phenyl ester labeling of RNA**. In a sterile 200 µL PCR tube, 2.7 µL 18nt test ssRNA (280 µM stock) was combined with 5.0 µL acetonitrile (to aid reagent solubility), 3.0 µL DMAP stock (400 mM in 50% aqueous DMSO), and 3.0 µL Coumarin Phenyl Ester stock/suspension in DMSO (200 mM). (Note: the coumarin phenyl ester stock was not fully soluble at 200 mM in

DMSO, but was sonicated briefly prior to pipetting to facilitate suspension and small particle size). The mixture was incubated for 6 h at 23°C with occasional vortexing. The mixture was then subjected to ethanol precipitation twice to isolate the RNA and remove unreacted Coumarin-PE reagent. The obtained pellet was then dissolved in 15 µL nuclease-free water for analysis by mass spectrometry, PAGE gel, and fluorescence spectroscopy.

**RNA acetylation with acetyl-CoA**. In a sterile PCR tube, 3 µL of 3.3x pH 7.5 MOPS buffer (333 mM MOPS, pH 7.5; 333 mM NaCl, 20 mM $MgCl_2$) was mixed with 1 µL of a 100 µM stock solution of an 18-nt ssRNA or ssDNA. Acetyl-CoA (100 mM) and DMAP (500 mM) were prepared as 5x stock solutions in water. 2 µL of each 5x stock was added to the reaction to achieve final concentrations of 20 mM Acetyl-CoA and 100 mM DMAP. The reaction was then brought to a final volume of 10 µL with nuclease-free water. Reactions were incubated at 23 °C for 24 h, followed by purification via ethanol precipitation. The extent of RNA modification was analyzed using quantitative MALDI-TOF MS.

**PAGE analysis of RNA fluorescent labeling.** 5 picomoles of Coumarin-PE-treated 18 nt RNA or unmodified RNA were mixed with 5 µL of 2X RNA Loading Dye and loaded onto a 12% polyacrylamide gel. Electrophoresis was performed in 1X TBE buffer (pH 8.3, Sigma Aldrich) at 15 W for 1 hr. Labeled RNA was visualized by fluorescence imaging using a coumarin filter. The gel was subsequently stained with SYBR Gold and re-imaged to detect all RNA species. Image analysis was performed using ImageJ/FIJI.

**Fluorimeter Measurements**: 25 picomoles of Coumarin-PE-treated 18 nt RNA (or unmodified RNA) was dissolved in 500 µL 1X PBS and transferred to a quartz cuvette. Using a Horiba Jobin-Yvon Spex Fluorolog-3™ fluorimeter and the FluoroEssence™ software, emission spectra were recorded with excitation at 420 nm, emission range 400-800 nm and slit width 5 nm.

**Measurements of reagent sensitivity to moisture:** Stability of electrophilic reagents in water was measured by analyzing changes in $^1$H NMR spectra over time during incubation at room temperature (23°C). Each reagent was dissolved at 5 mM in *ca*. 700 µL $D_2O$ with 15% v/v DMSO-d6 (2-chloropyrimidine-4-carbonitrile, phenyl thioacetate, phenyl acetate) or 30% (benzoic anhydride, methyl imidazole carbamate, benzenesulfonyl imidazole) to ensure solubility. Initial rate measurements were employed to minimize nonlinearity due to effects of pH change during reactions. $^1$H-NMR spectra were recorded on a Varian Mercury 500 MHz instrument at different time intervals and reaction progress was monitored from the relative peak intensities of reagents and hydrolysis products. The rate constants and half-lives of hydrolysis were determined by fitting the obtained data to the first-order rate equation. Line fits are shown in Fig. S8.

**MALDI-TOF mass spectrometry:** All MALDI-TOF spectra were recorded at the Stanford University Mass Spectrometry facility, using the Bruker Daltonik Microflex MALDI-TOF spectrometer equipped with an $N_2$ laser. Spectra were recorded in linear negative mode and samples were plated on an MSP Anchorchip 96 target plate. 0.3 M trihydroxyacetophenone in EtOH (matrix) and 0.1 M aqueous ammonium citrate (co-matrix) were mixed in a 4:1 ratio by volume to be used as a matrix mix for MALDI. This mix was freshly prepared before analysis.

After RNA precipitation, the RNA pellet was redissolved in 2.5 µL RNAse-free water to prepare a ~60 µM sample solution. 0.8 µL of the matrix mix and 0.8 µL of the RNA solution were spotted successively on the plate and allowed to air dry 15 min. The spectral data was then recorded using Flex Control software (Bruker), and analyzed using MNova software (Mestrenova).

**NMR Spectroscopy:** All NMR spectra were recorded at the Stanford University Department of Chemistry NMR facility. Varian 300 MHz, 400 MHz and 500 MHz NMR instruments were used to record the $^{1}H$ and $^{13}C$ spectra. The spectra were analyzed using MNova software.

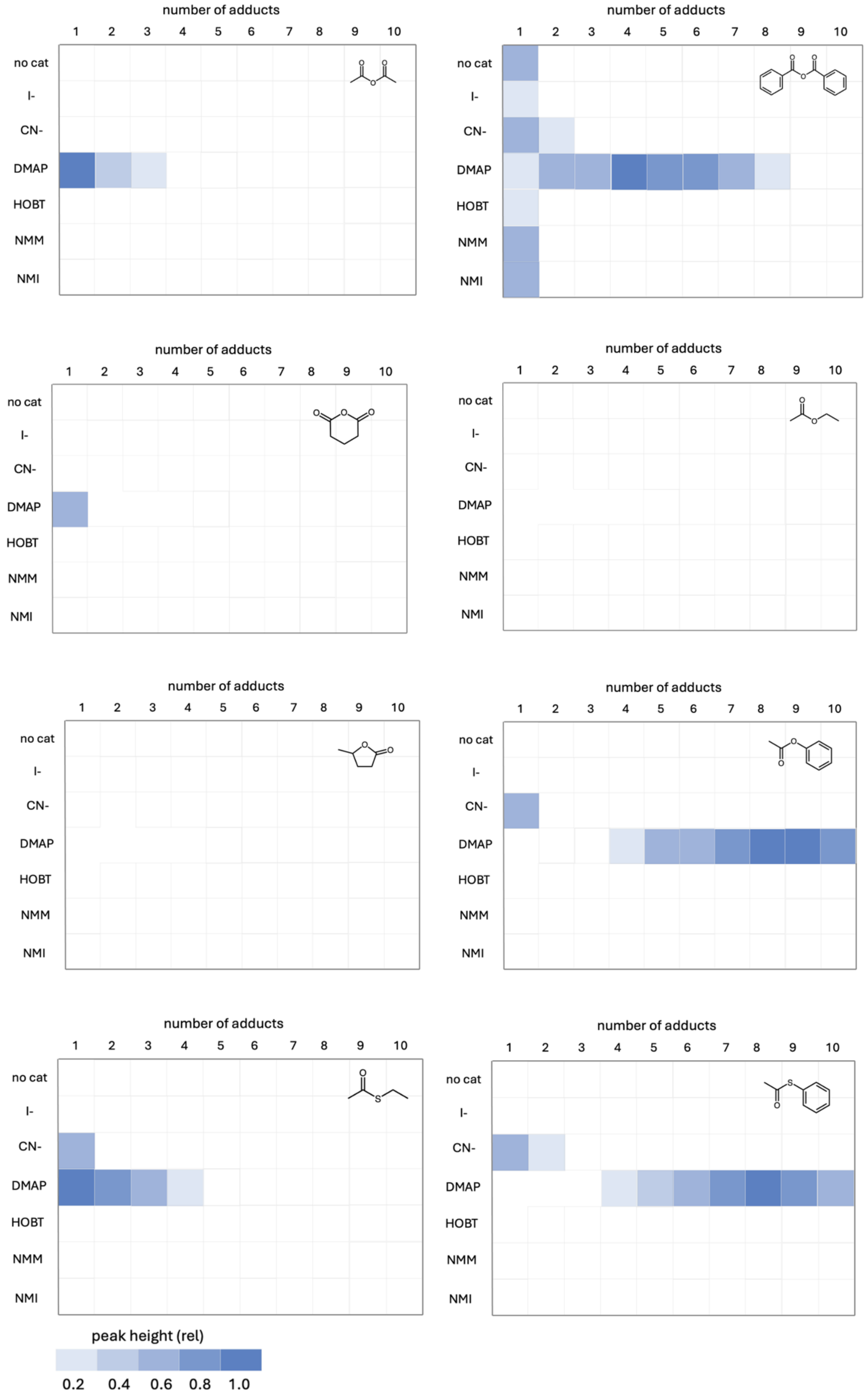
number of adducts
1 2 3 4 5 6 7 8 9 10
no cat
I-
CN-
DMAP
HOBT
NMM
NMI
peak height (rel)
0.2 0.4 0.6 0.8 1.0

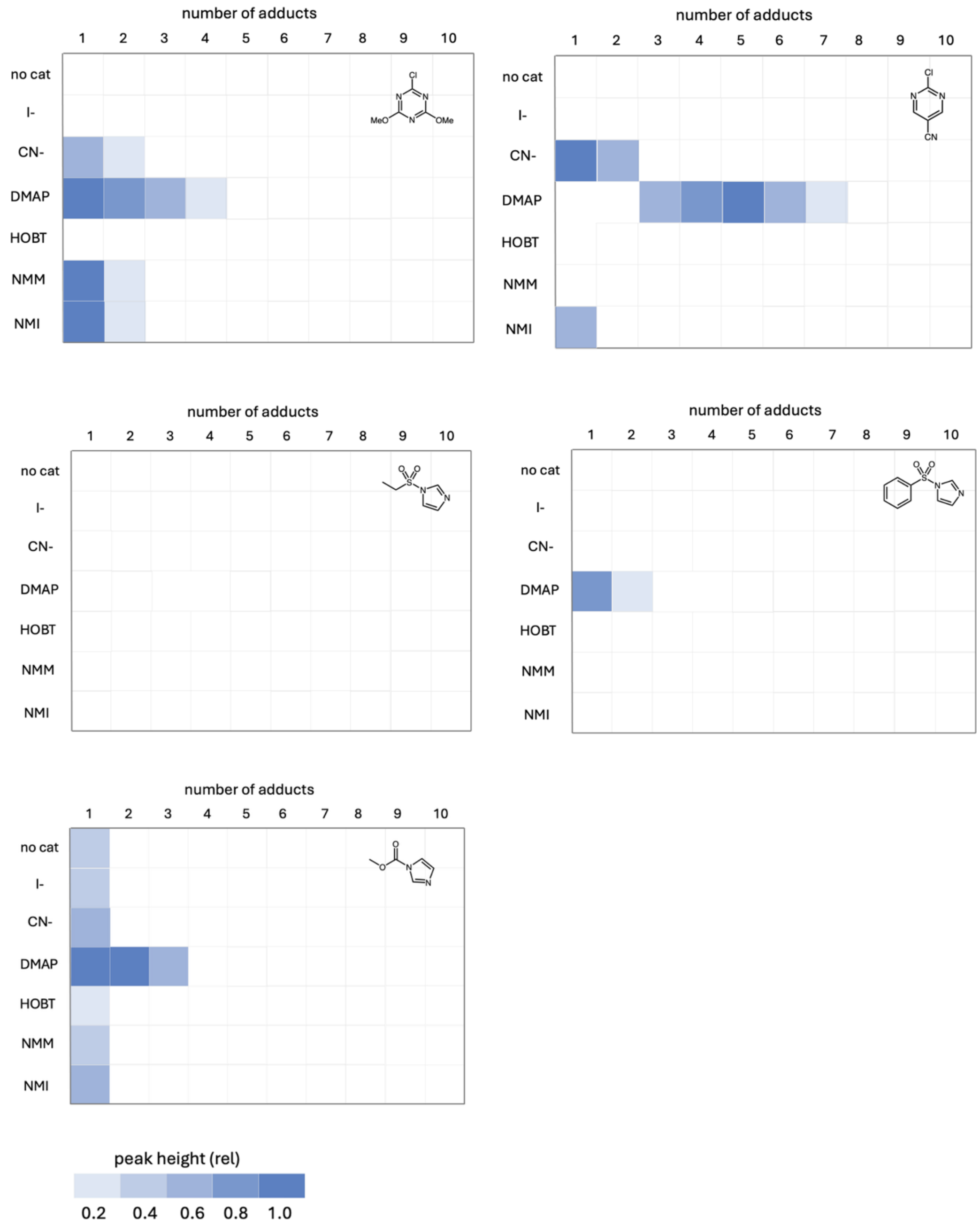

**Figure S1**. Heat maps of reactivity of 13 electrophilic reagents (structures shown) with single-stranded RNA (9 uM) in the presence of 6 different nucleophilic species as potential catalysts. Color scale of mass spectrometric peak heights is shown. Reactions were performed in phosphate-buffered saline (pH 7.4) supplemented to 30 mM phosphate (23°C, 6 h). Yields were determined by quantitative MALDI-TOF mass spectrometry as described above. Experiments were performed in duplicate and results averaged.

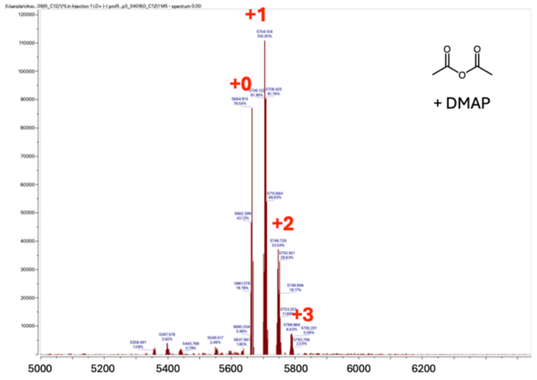
+1
+0
+2
+3
+ DMAP
5000
5200
5400
5600
5800
6000
6200

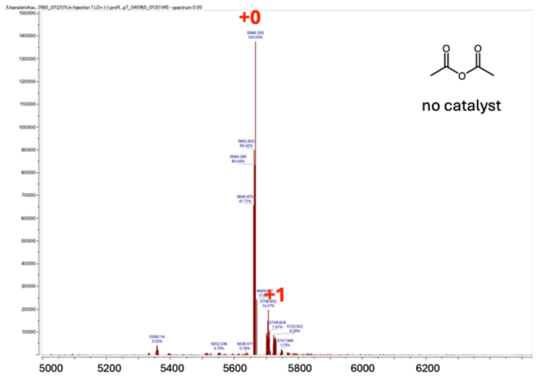
+0
+1
no catalyst
5000
5200
5400
5600
5800
6000
6200

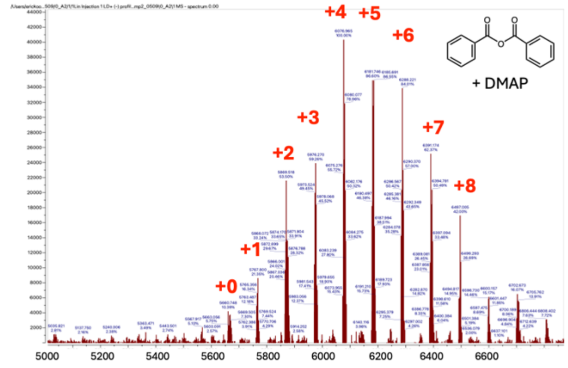
+4
+5
+6
+3
+2
+7
+8
+1
+0
+ DMAP
5000
5200
5400
5600
5800
6000
6200
6400
6600

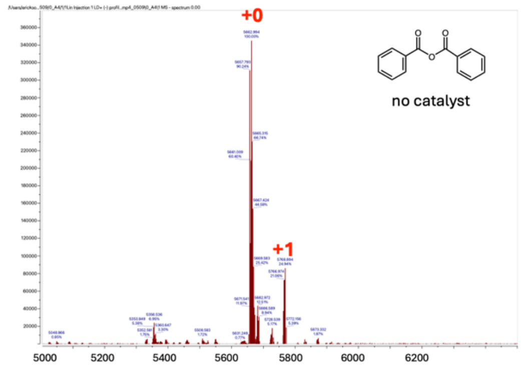
+0
+1
no catalyst
5000
5200
5400
5600
5800
6000
6200

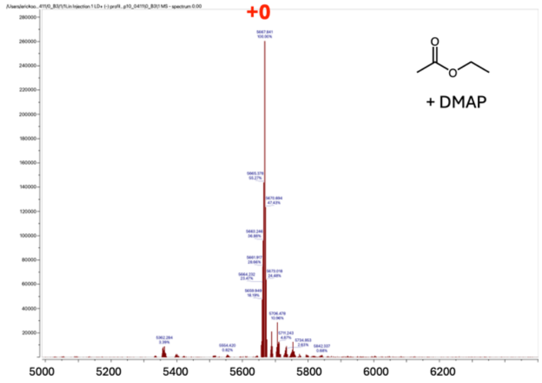
+0
+ DMAP
5000
5200
5400
5600
5800
6000
6200

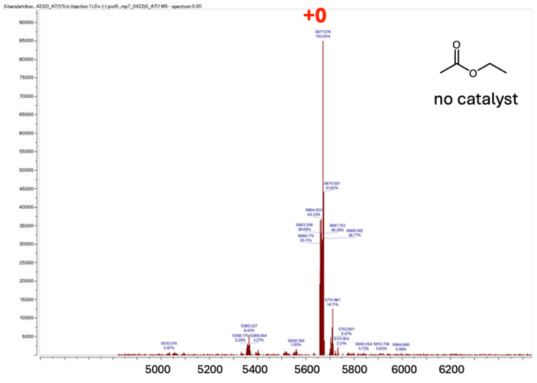
+0
no catalyst
5000
5200
5400
5600
5800
6000
6200

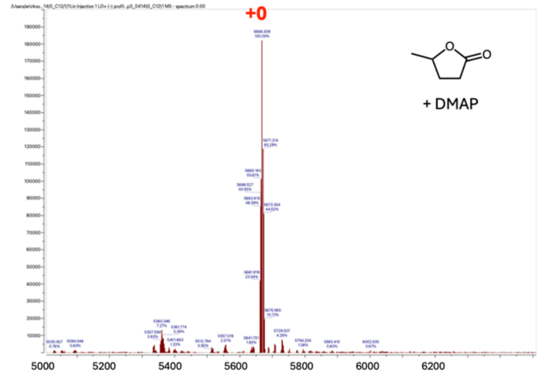
+0
+ DMAP
5000
5200
5400
5600
5800
6000
6200

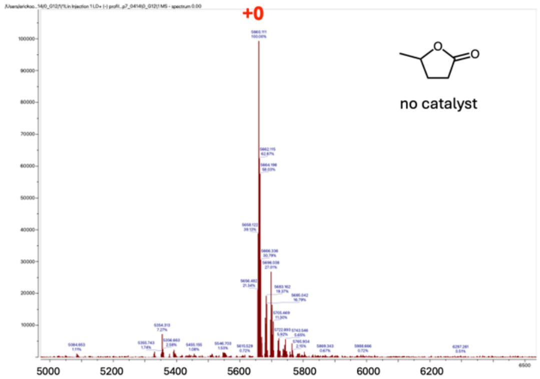
+0
no catalyst
5000
5200
5400
5600
5800
6000
6200

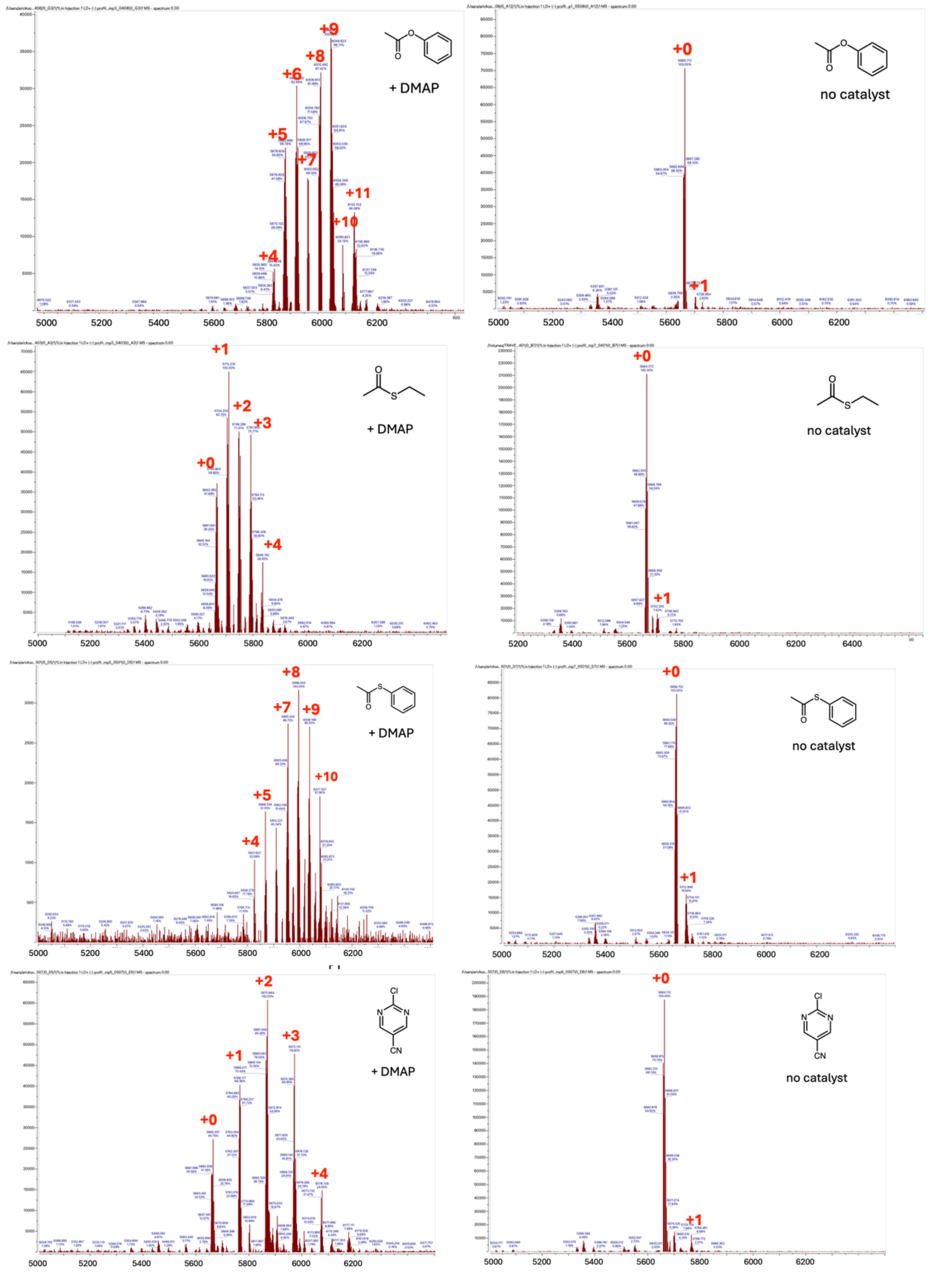
+9
+8
+6
+5
+7
+11
+10
+4
+ DMAP
+0
+1
no catalyst
+1
+2
+3
+0
+4
+ DMAP
+0
+1
no catalyst
+8
+7
+9
+10
+5
+4
+ DMAP
+0
+1
no catalyst
+2
+3
+1
+0
+4
Cl
N
N
CN
+ DMAP
+0
+1
Cl
N
N
CN
no catalyst

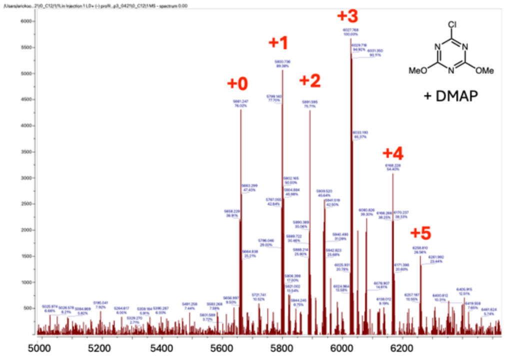
+0
+1
+2
+3
+4
+5
MeO
OMe
Cl
N
+ DMAP
5000
5200
5400
5600
5800
6000
6200

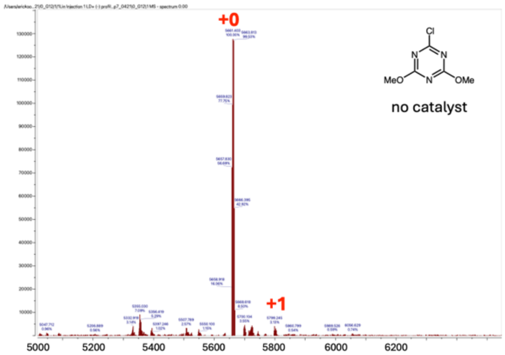
+0
+1
MeO
OMe
Cl
N
no catalyst
5000
5200
5400
5600
5800
6000
6200

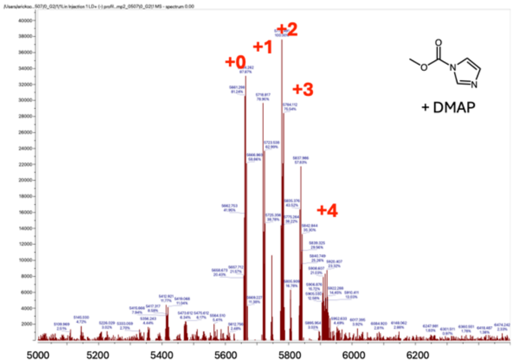
+0
+1
+2
+3
+4
+ DMAP
5000
5200
5400
5600
5800
6000
6200

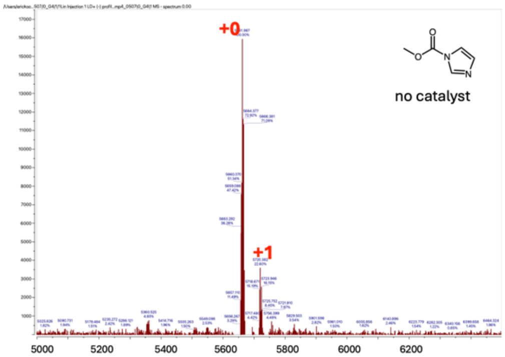
+0
+1
no catalyst
5000
5200
5400
5600
5800
6000
6200

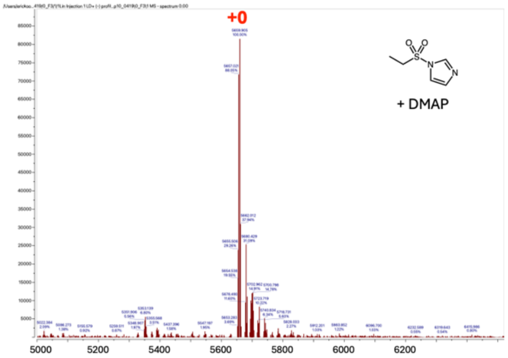
+0
+ DMAP
5000
5200
5400
5600
5800
6000
6200

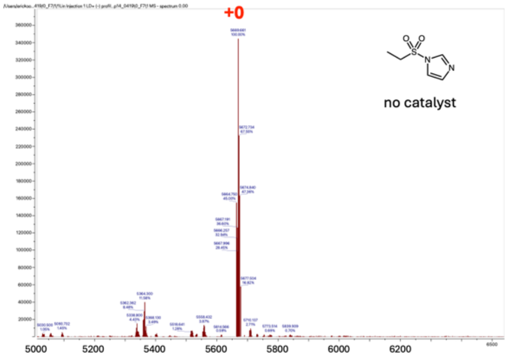
+0
no catalyst
5000
5200
5400
5600
5800
6000
6200

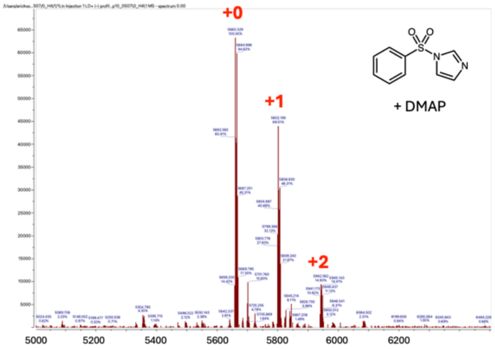
+0
+1
+2
+ DMAP
5000
5200
5400
5600
5800
6000
6200

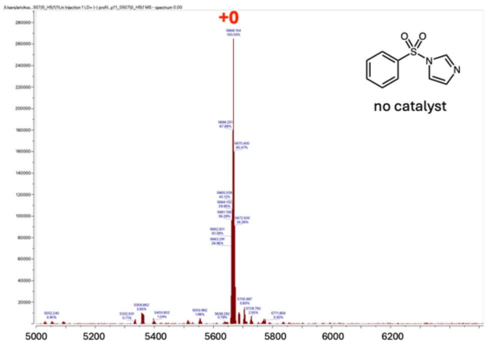
+0
no catalyst
5000
5200
5400
5600
5800
6000
6200

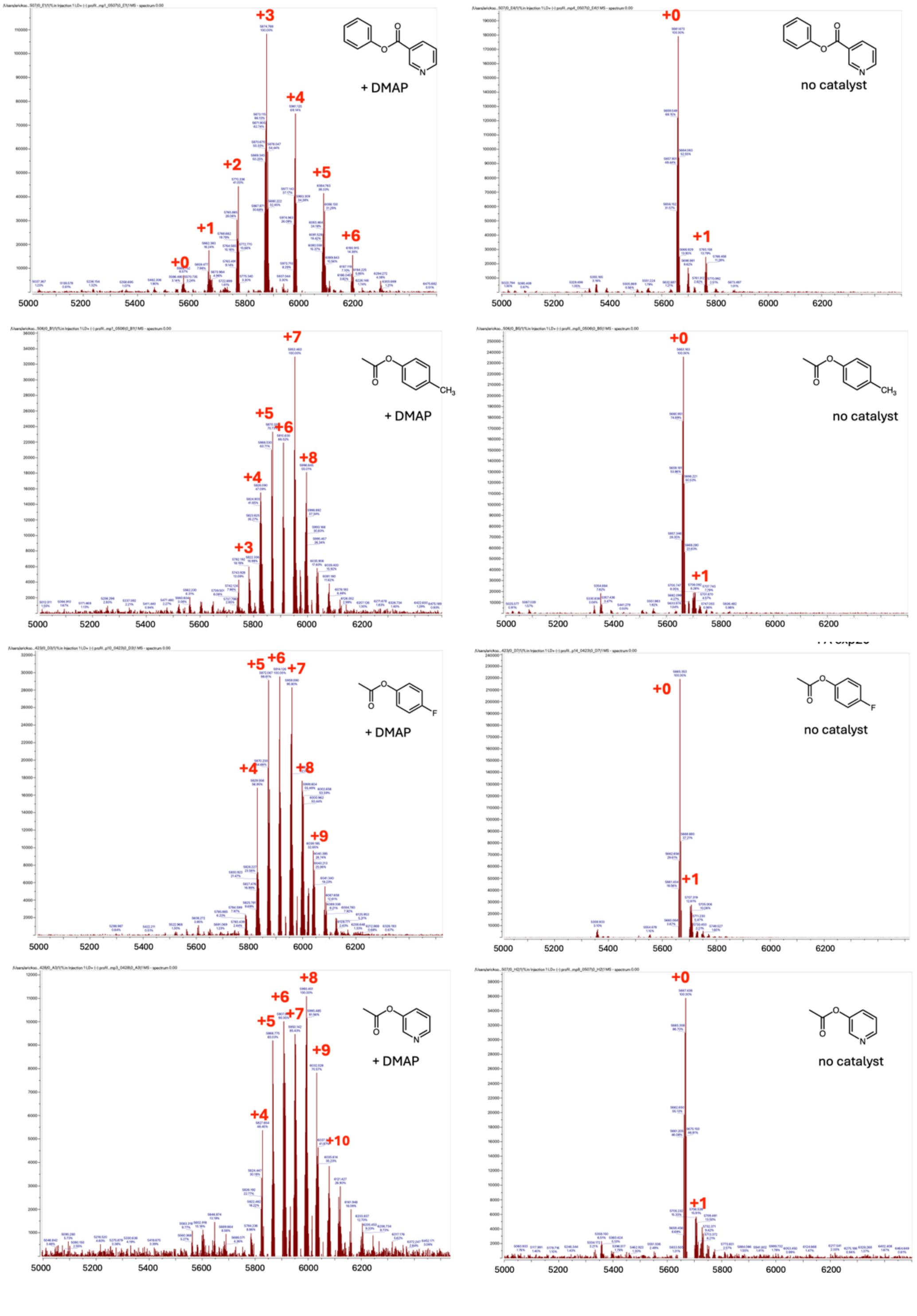

**Figure S2**. Representative MALDI-TOF mass spectrometry raw data documenting reactivity of electrophilic reagents with 9 μM test ssRNA (sequence 5′AUCCUGCCGACUACGCCA) in the presence and absence of DMAP (20 mM) (“no cat”). Reagents (40 mM ) were tested at 23 °C for 6 h in pH 7.4 phosphate-buffered saline (PBS) buffer supplemented to 30 mM phosphate, containing 15% DMSO. Red numerals indicate number of adducts per RNA strand (MW=5650 Da).

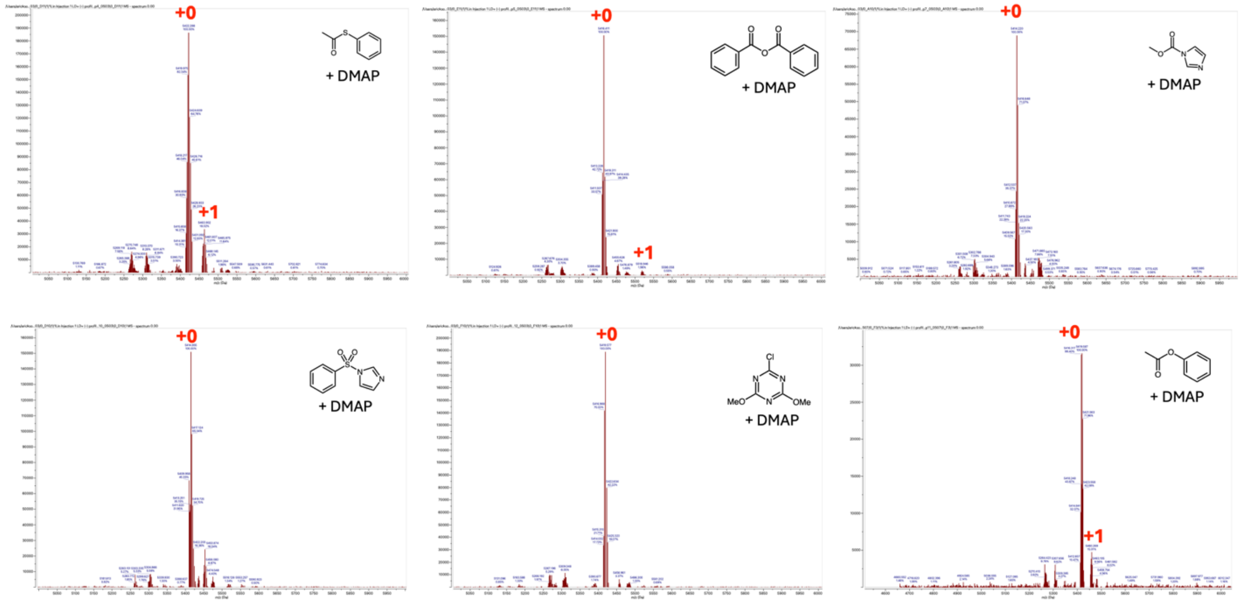

**Figure S3**. MALDI-TOF spectra of control reactions showing low reactivity of DMAP-responsive reagents with single-stranded DNA. Conditions were as for Fig. S2, with 20 mM DMAP. The spectra document little or no adduct formation with ssDNA, providing evidence that the reactions with RNA (see Fig. 2) occur largely or completely at 2′-OH groups rather than exocyclic amine groups of the nucleobases. Red numerals above peaks indicate number of adducts on the DNA strand (MW of unmodified DNA = 5404 Da).

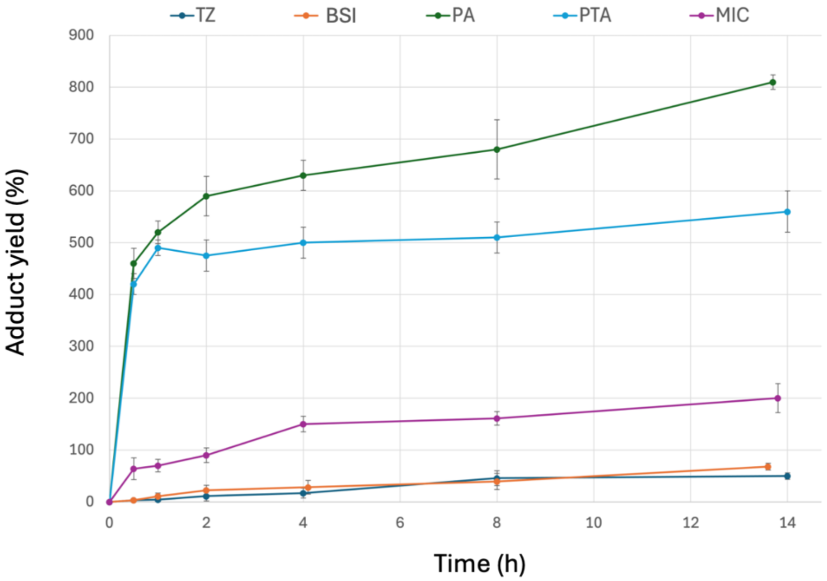

**Figure S4**. Full time course of RNA reaction with five classes of reagents (40 mM) in the presence of DMAP (20 mM). Abbreviations: TZ=chloro-dimethoxytetrazine; BSI=benzenesulfonyl imidazole; PA=phenyl acetate; PT=phenyl thioacetate; MIC=methyl imidazole carbamate. Details are given in the main text, Fig. 3. Experiments were performed in quadruplicate and results averaged; error bars show standard deviations.

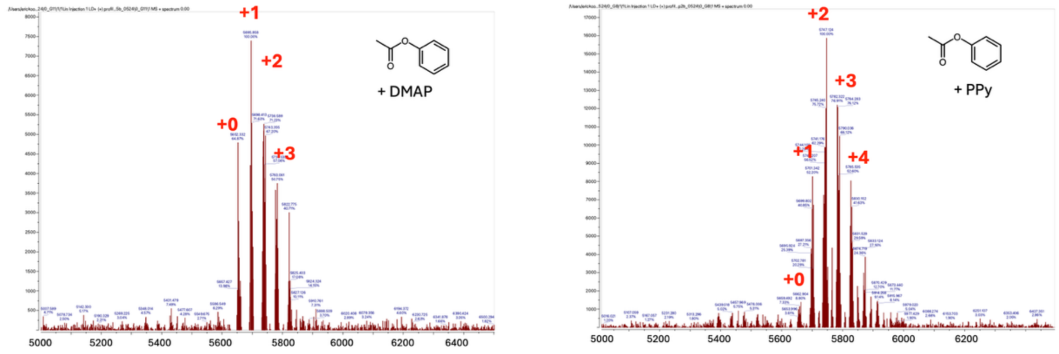

**Figure S5**. Representative mass spectrometry data comparing acetylation of ssRNA with phenyl acetate (40 mM) in the presence of DMAP or PPy (10 mM). Reactions were performed at 23 °C for 2 h in pH 7.4 phosphate-buffered saline (PBS) buffer supplemented to 30 mM phosphate, containing 15% DMSO. Experiments were carried out in quadruplicate, and average yields are reported in the main text.

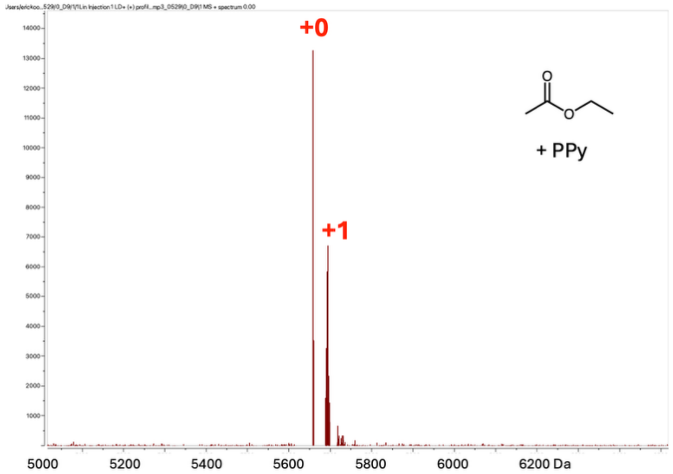

**Figure S6**. More active nucleophilic catalyst PPy enables significant RNA acylation by the poorly reactive donor ethyl acetate. Monoacetylated RNA yield (+1 peak) is ~33%. Conditions: 40 mM EtOAc, 40 mM PPy, RNA 9 μM, 23 °C, 6 h in pH 7.4 phosphate-buffered saline (PBS) buffer supplemented to 30 mM phosphate, containing 15% DMSO.

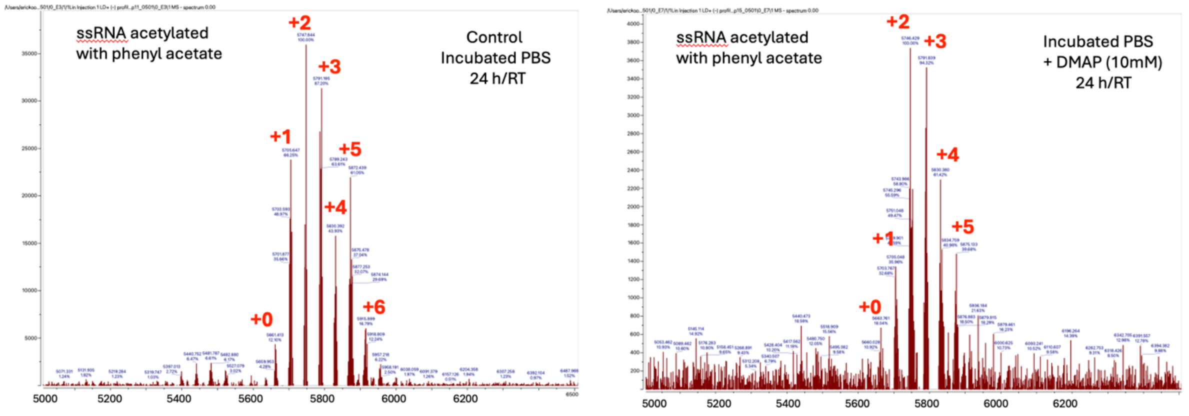

**Figure S7**. Experiments testing the ability of DMAP to catalyze loss of acetyl groups from RNA in the absence of acetylating reagent. (left) example of MALDI-TOF spectrum of acetylated RNA incubated in pH 7.4 buffer (24 h, 23 °C).; (right) same acetylated RNA incubated in buffer with 10 mM DMAP under identical conditions. Results show only modest loss of acetyl groups under these conditions. Experiments were performed in quadruplicate; average number of acetyl groups in the control group was 2.8 +/- 0.8; average for DMAP-added samples was 2.1 +/- 1.5.

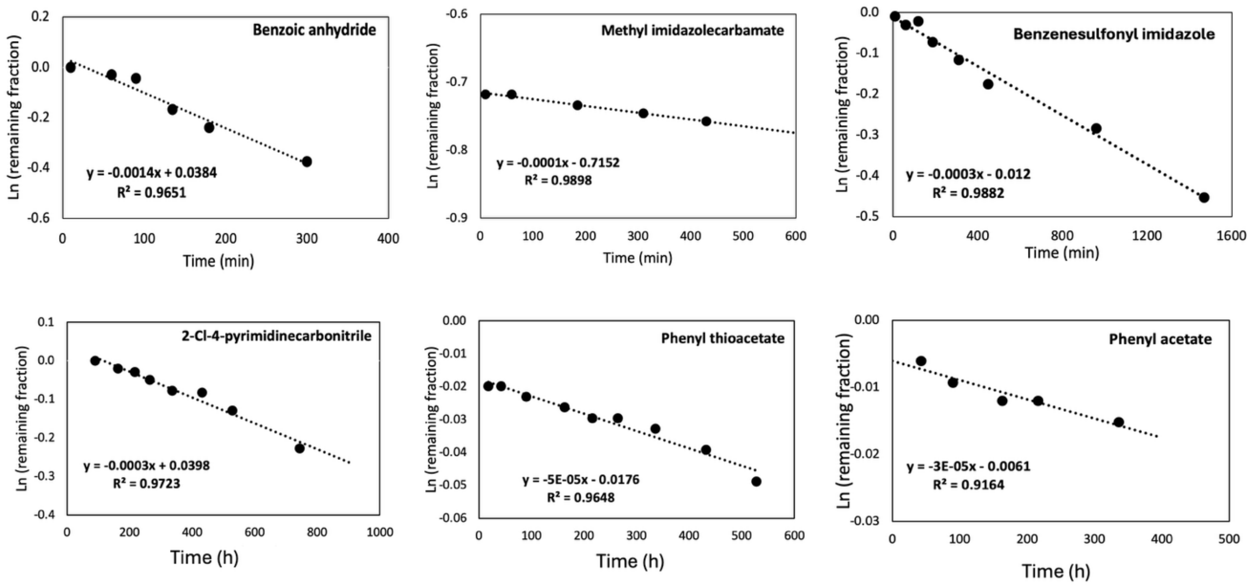

**Figure S8**. Kinetics plots for measurement of aqueous half-lives for six RNA-modifying reagents by $^{1}H$ NMR. Shown are plots of hydrolysis data for selected classes of reagents in $D_2O$. Conditions: 5 mM reagent, 23°C, $D_2O$ with 15% DMSO-d6 (top row) or 30% DMSO-d6 (bottom row).

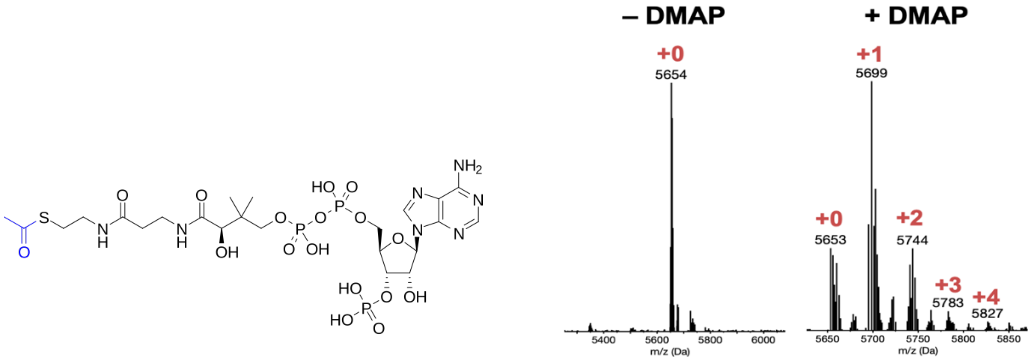

**Figure S9**. DMAP-assisted RNA acetylation with Acetyl CoA. The biological thioester acetyl donor (structure shown) was incubated with an 18nt RNA at pH 7.5 over 24h in the absence and presence of DMAP (100 mM) (see Methods). Results confirmed up to 4 acetyl groups transferred.

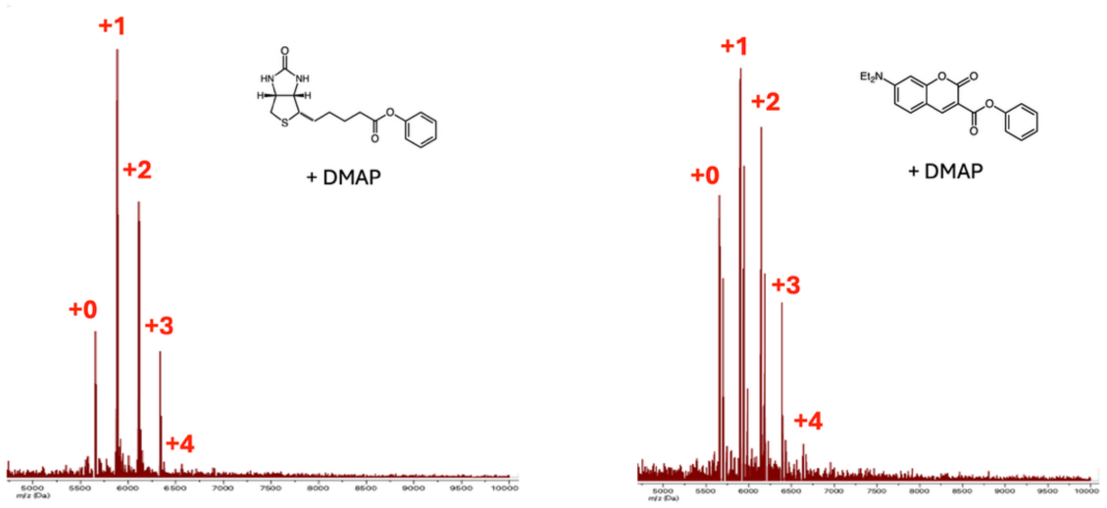

**Figure S10**. Mass spectrometry data documenting DMAP-assisted labelling of 18 nt ssRNA with Biotin-PE (left) and fluorescent Coumarin-PE reagent (right) after 6h reaction. RNA conversion yields were 85% and 79% respectively. Red numerals indicate number of adducts per RNA strand (MW=5650 Da).

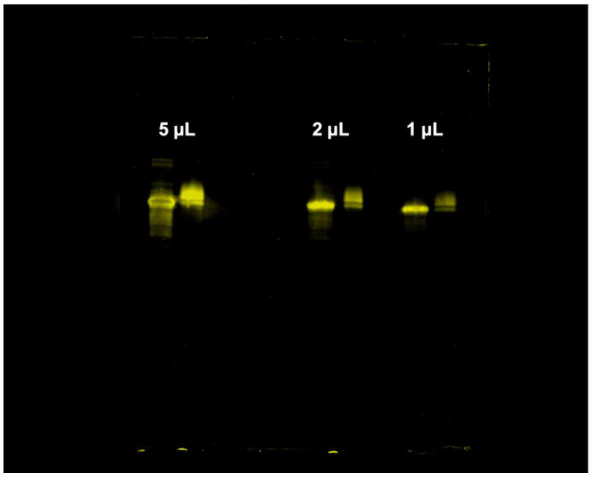

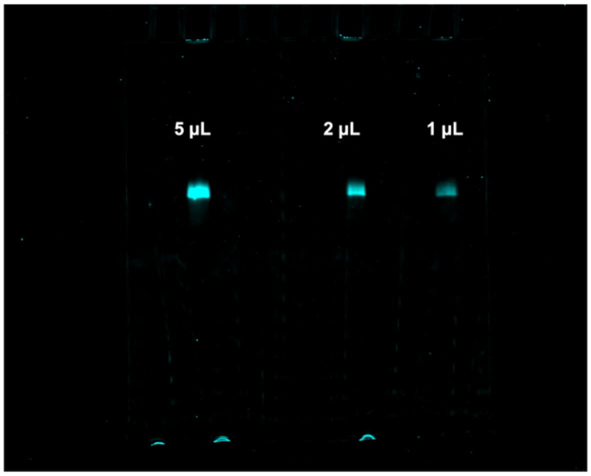

**Figure S11**. Fluorescence images of full uncropped PAGE gel of 18 nt RNA labeled with Coumarin-PE. Three lane groups show RNA loaded in decreasing amounts (5 µL = 25 pmole); yellow image is image taken with SYBR Gold filter; cyan image is taken with coumarin filter. Left lane of each group is unmodified RNA; note that the coumarin dye slows gel mobility slightly.

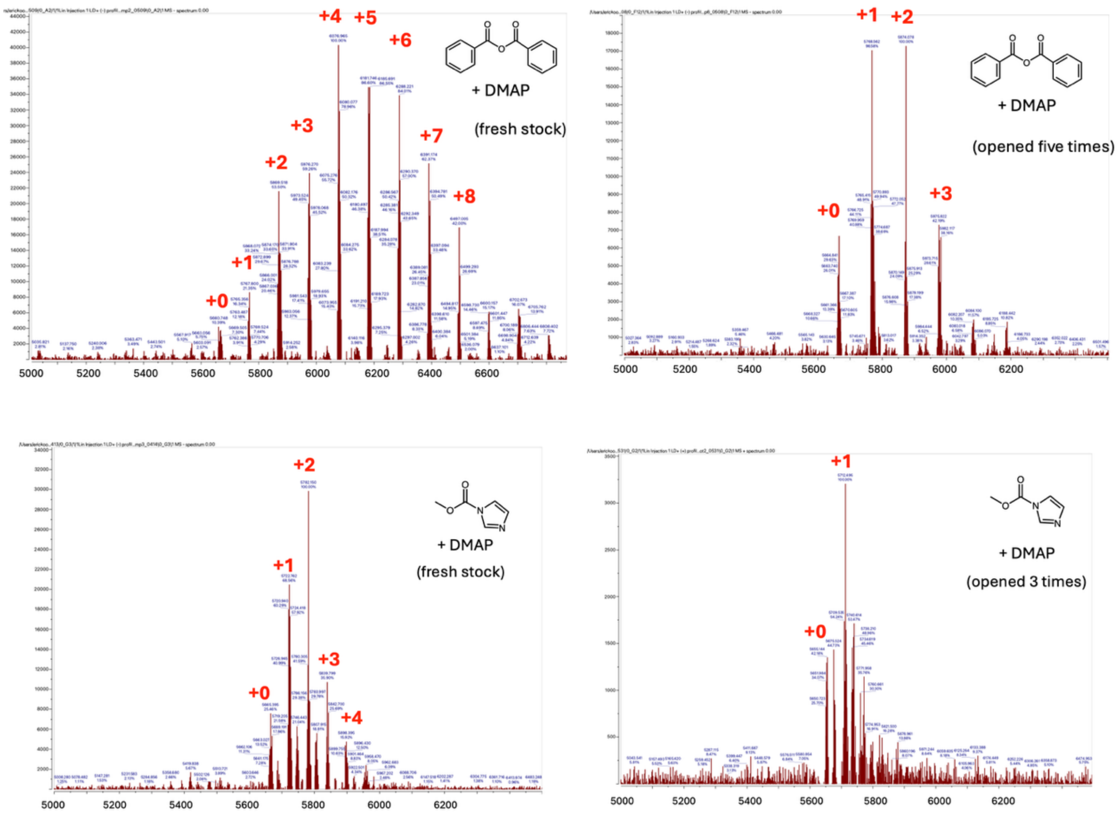

**Figure S12**. Loss of RNA acylation activity of benzoic anhydride and methyl imidazolecarbamate stock solutions after being opened, as evidenced by reduction in number of acyl adducts seen by mass spectrometry. Reactions were performed under identical conditions (see Fig. 2 in main text). Original stock solutions were prepared in dry DMSO. Stocks were stored frozen at -20°, and were warmed to room temperature before briefly opening for each use.

**Synthesis of reagents**

**1.** Synthesis of 1-(benzenesulfonyl)-1H-imidazole

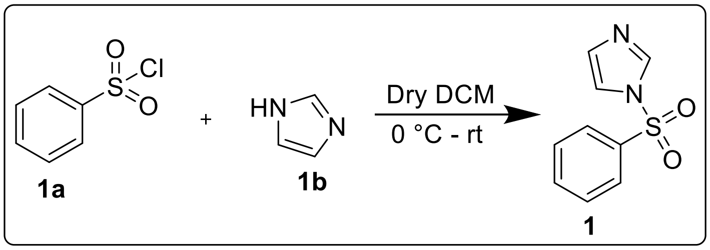

Compound **1** was synthesized using the method described in reference 1. To a stirred solution of imidazole (**1b**, 612 mg, 9 mmol) in dry dichloromethane (20 mL) at 0 °C, benzenesulfonyl chloride (**1a**, 530 mg, 3 mmol) was added dropwise using a syringe. The reaction mixture was allowed to stir at room temperature for 5 hours. Upon completion of the reaction as indicated by TLC, the reaction mixture was washed with water (2x) followed by brine (2x). The organic solvent was distilled off and the crude residue was subjected to column chromatography using silica gel (60–120 mesh) as the stationary phase and EtOAc in hexane (0 percent to 20 percent) as the mobile phase to afford the title compound.

White solid, Yield 96% $^1$H NMR (500 MHz, DMSO-$d_6$) δ 8.39 (t, J = 1.1 Hz, 1H), 8.12 – 8.08 (m, 2H), 7.85 – 7.81 (m, 1H), 7.77 (d, J = 1.5 Hz, 1H), 7.74 – 7.69 (m, 2H), 7.14 (dd, J = 1.6, 0.9 Hz, 1H). $^{13}$C NMR (125 MHz, DMSO-$d_6$) δ 137.8, 135.9, 131.8, 130.7, 127.7, 118.8. HRMS (ESI) m/z [M + H]$^+$: calcd for $C_9H_9N_2O_2S$: 209.0385; found: 209.0377.

**2.** Synthesis of 1-(ethanesulfonyl)-1H-imidazole

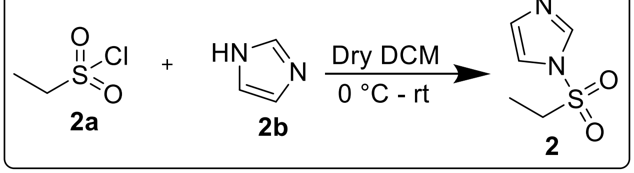

Compound **2** was synthesized using the method described in reference 1. To a stirred solution of imidazole (**2b**, 817 mg, 12 mmol) in dry dichloromethane (20 mL) at 0 °C, ethanesulfonyl chloride (**2a**, 514 mg, 4 mmol) was added dropwise using a syringe. The reaction mixture was allowed to stir at room temperature for 5 hours. Upon completion of the reaction as indicated by TLC, the reaction mixture was washed with water (2x) followed by brine (2x). The organic solvent was distilled off and the crude residue was subjected to column chromatography using silica gel (60–120 mesh) as the stationary phase and methanol in DCM (0 percent to 2 percent) as the mobile phase to afford the title compound.

White solid, Yield 99% $^{1}$H NMR (500 MHz, DMSO-$d_6$) δ 8.18 – 8.17 (m, 1H), 7.65 (t, J = 1.4 Hz, 1H), 7.19 (dd, J = 1.6, 0.9 Hz, 1H), 3.77 (q, J = 7.3 Hz, 2H), 1.16 – 1.12 (m, 3H). $^{13}$C NMR (125 MHz, DMSO-$d_6$) δ 137.8, 131.0, 119.1, 49.7, 8.1. HRMS (ESI) m/z [M + H]$^+$: calcd for $C_5H_9N_2O_2S$: 161.0385; found: 161.0377.

**3.** Synthesis of 3-hydroxypyridinyl acetate

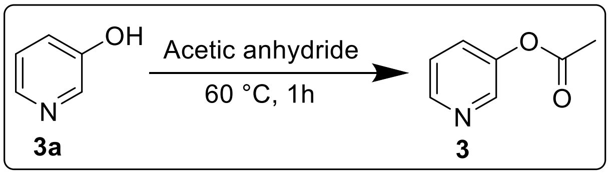

Compound **3** was synthesized using the method described in reference 2. A stirred solution of 3-hydroxypyridine (**3a**, 951 mg, 10 mmol) in acetic anhydride (5.1 g, 50 mmol) was heated at 60 °C for 1 hour. Upon completion of the reaction as indicated by TLC, 50 mL of dry diethyl ether was added. The organic layer was washed with $NaHCO_3$ (2x) followed by water (2x) and dried over $MgSO_4$. The organic solvent was distilled off and the crude residue was subjected to column chromatography using silica gel (60–120 mesh) as the stationary phase and ethyl acetate in hexane (0 percent to 50 percent) as the mobile phase to afford the title compound.

Yellow liquid, Yield 88% $^{1}$H NMR (600 MHz, DMSO-$d_6$) δ 8.43 (dd, J = 4.7, 1.4 Hz, 1H), 8.38 (d, J = 2.7 Hz, 1H), 7.59 (ddd, J = 8.2, 2.7, 1.4 Hz, 1H), 7.45 – 7.42 (m, 1H), 2.27 (s, 3H). $^{13}$C NMR (150 MHz, DMSO-$d_6$) δ 169.5, 147.6, 147.3, 143.8, 130.1, 124.8, 21.2. HRMS (ESI) m/z [M + H]$^+$: calcd for $C_7H_8NO_2$: 138.0555; found: 138.0548.

**4.** Synthesis of biotin phenyl ester

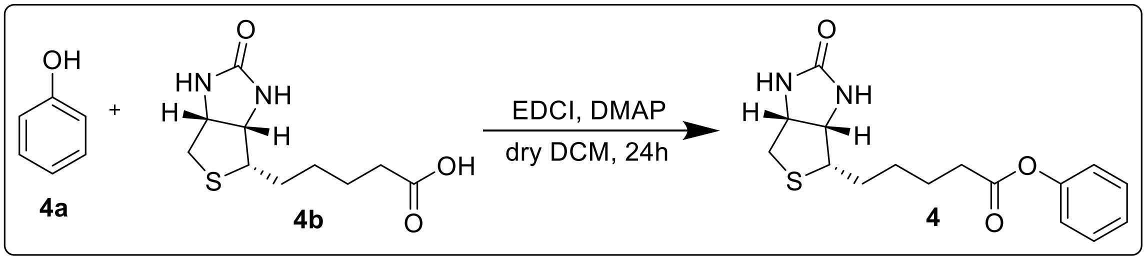

Compound **4** was synthesized using the method described in reference 3. To a stirred solution of phenol (**4a**, 94 mg, 1 mmol) and D-Biotin (**4b**, 269 mg, 1.1 mmol) in dry DCM (20 mL) were added EDCI (480 mg, 2.5 mmol) and DMAP (80 mg, 0.65 mmol). The resulting solution was stirred at room temperature for 24 hours. Upon completion of the reaction as indicated by TLC, water (20 mL) was added to quench the reaction. The organic layer was separated and distilled off to afford a crude residue which was subjected to column chromatography using silica gel (60–120 mesh) as the stationary phase and methanol in DCM (0 percent to 4 percent) as the mobile phase to afford the title compound.

White solid, Yield 78% $^1$H NMR (500 MHz, DMSO-$d_6$) δ 7.45 – 7.40 (m, 2H), 7.26 (t, J = 7.4 Hz, 1H), 7.12 (dt, J = 7.8, 1.1 Hz, 2H), 6.44 (s, 1H), 6.36 (s, 1H) 4.32 (dd, J = 7.7, 5.2 Hz, 1H), 4.19 – 4.14 (m, 1H), 3.19 – 3.12 (m, 1H), 2.85 (dd, J = 12.4, 5.1 Hz, 1H), 2.64 – 2.56 (m, 3H), 1.74 – 1.64 (m, 3H), 1.58 – 1.40 (m, 3H). $^{13}$C NMR (125 MHz, DMSO-$d_6$) δ 172.2, 163.2, 151.0, 129.9, 126.2, 122.3, 61.5, 59.7, 55.8, 33.8, 28.5, 28.4, 24.9. HRMS (ESI) m/z $[M + H]^+$: calcd for $C_{16}H_{21}N_2O_3S$: 321.1273; found: 321.1264.

**5.** Synthesis of phenyl 7-(diethylamino)-2-oxo-2H-chromene-3-carboxylate

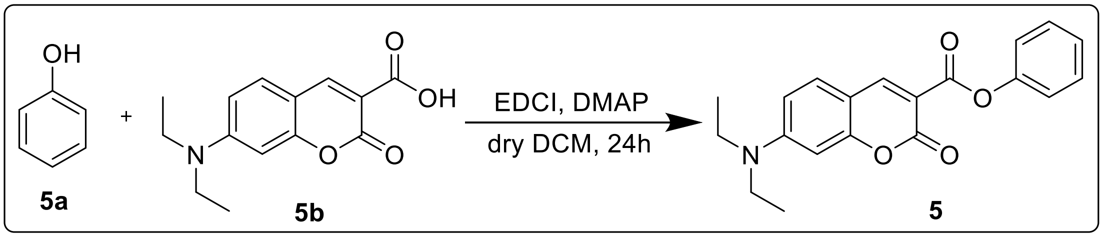

Compound **5** was synthesized using the method described in reference 3. To a stirred solution of phenol (**5a**, 16 mg, 0.17 mmol) and 7-(diethylamino)-2-oxo-2H-chromene-3-carboxylic acid (**5b**, 49 mg, 0.19 mmol) in dry DCM (3 mL) were added EDCI (82 mg, 0.43 mmol) and DMAP (14 mg, 0.11 mmol). The resulting solution was stirred at room temperature for 24 hours. Upon completion of the reaction as indicated by TLC, it was washed sequentially with water (5×2 mL), 1% acetic acid (5×2 mL) and water (5×2 mL). The organic layer was separated and distilled off to afford a yellow solid which was used without further purification.

Yellow solid, Yield 91% $^{1}$H NMR (500 MHz, DMSO-$d_6$) δ 8.82 (s,1H), 7.70 (d, J = 9.0 Hz, 1H), 7.50 – 7.44 (m, 2H), 7.30 (t, J = 7.3 Hz, 1H), 7.23 (d, J = 7.3 Hz, 2H), 6.83 (dd, J = 9.0, 2.5 Hz, 1H), 6.60 (d, J = 2.7 Hz, 1H), 3.52 (q, J = 7.1 Hz, 4H), 1.17 (t, J = 7.0 Hz, 6H). $^{13}$C NMR (125 MHz, DMSO-$d_6$) δ 162.33, 158.87, 157.44, 153.81, 151.15, 150.92, 132.64, 129.96, 126.20, 122.47, 110.55, 107.67, 106.25, 96.36, 44.92, 12.83. HRMS (ESI) m/z $[M + H]^+$: calcd for $C_{20}H_{20}NO_4$: 338.1392; found: 338.1386.

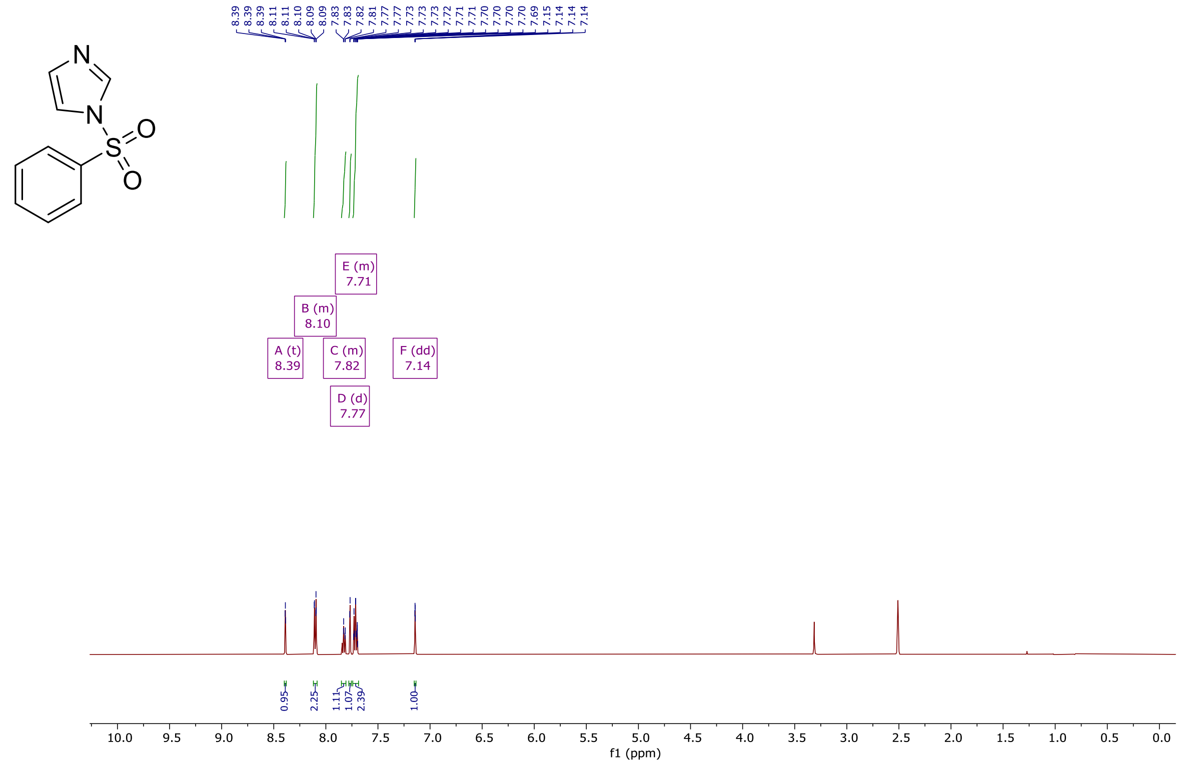

$^{1}$H NMR spectrum of compound **1** (500 MHz, DMSO-$d_6$)

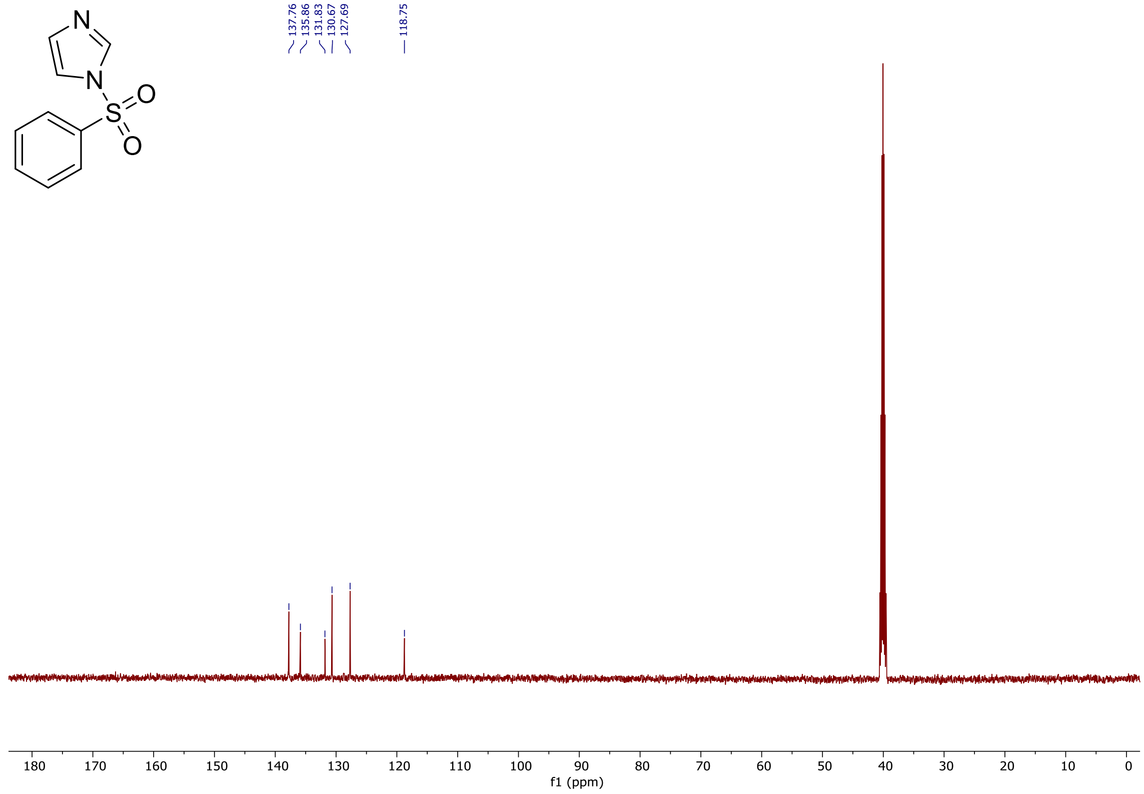

$^{13}$C NMR spectrum of compound **1**(125 MHz, DMSO-$d_6$)

tmcl_X240_35121.raw, C9H8N2O2S [H]+

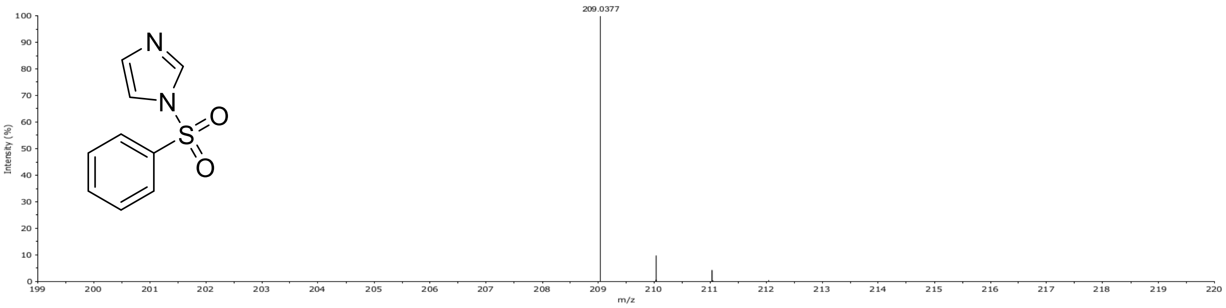

HRMS spectrum of compound **1**

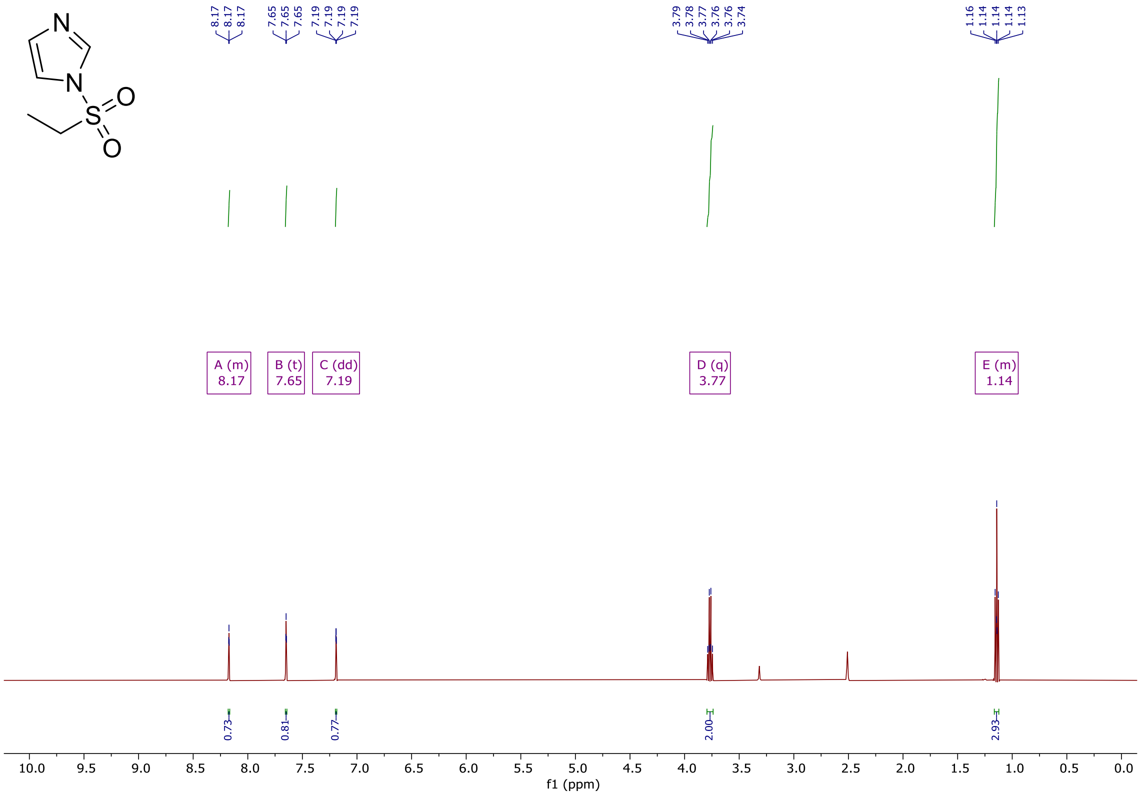

$^{1}$H NMR spectrum of compound **2** (500 MHz, DMSO-$d_6$)

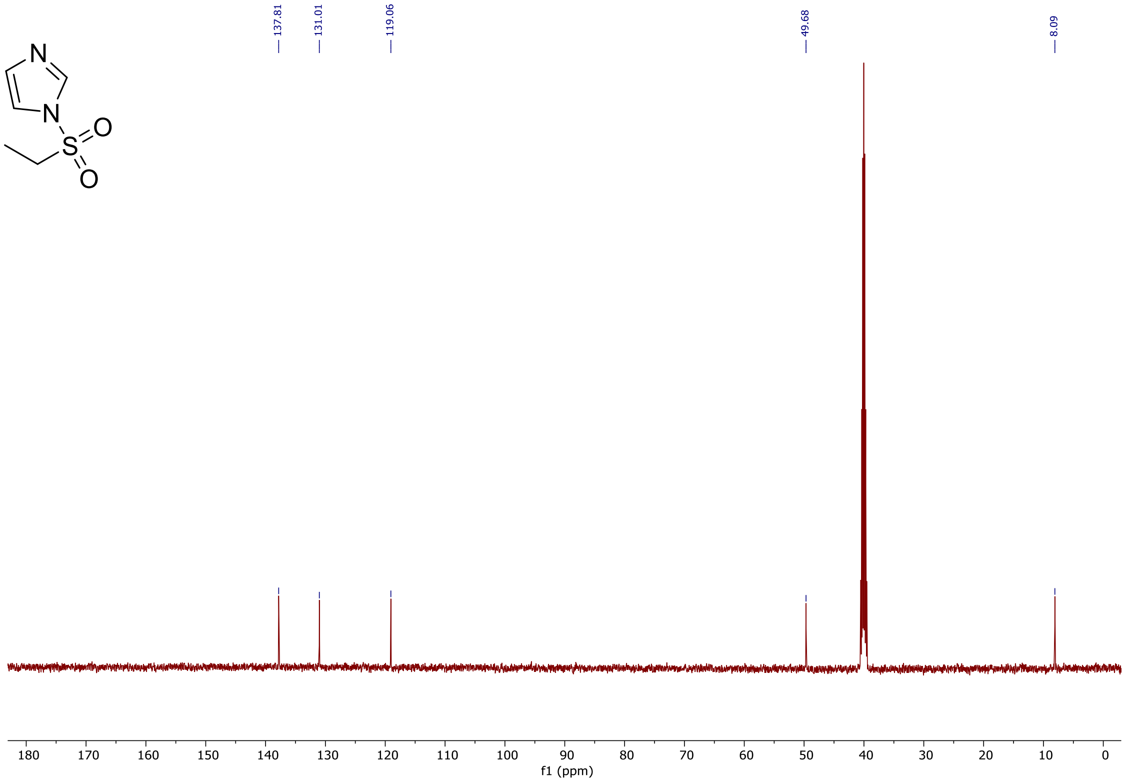

$^{13}$C NMR spectrum of compound **2** (125 MHz, DMSO-$d_6$)

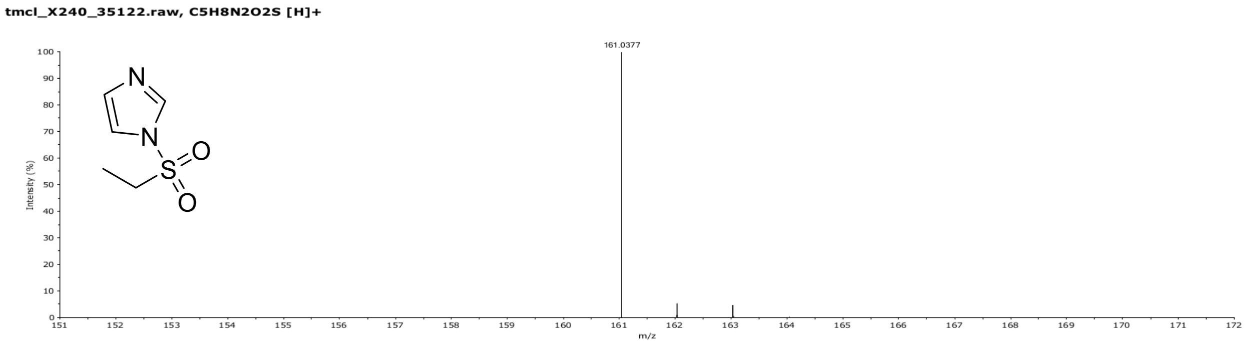

HRMS spectrum of compound **2**

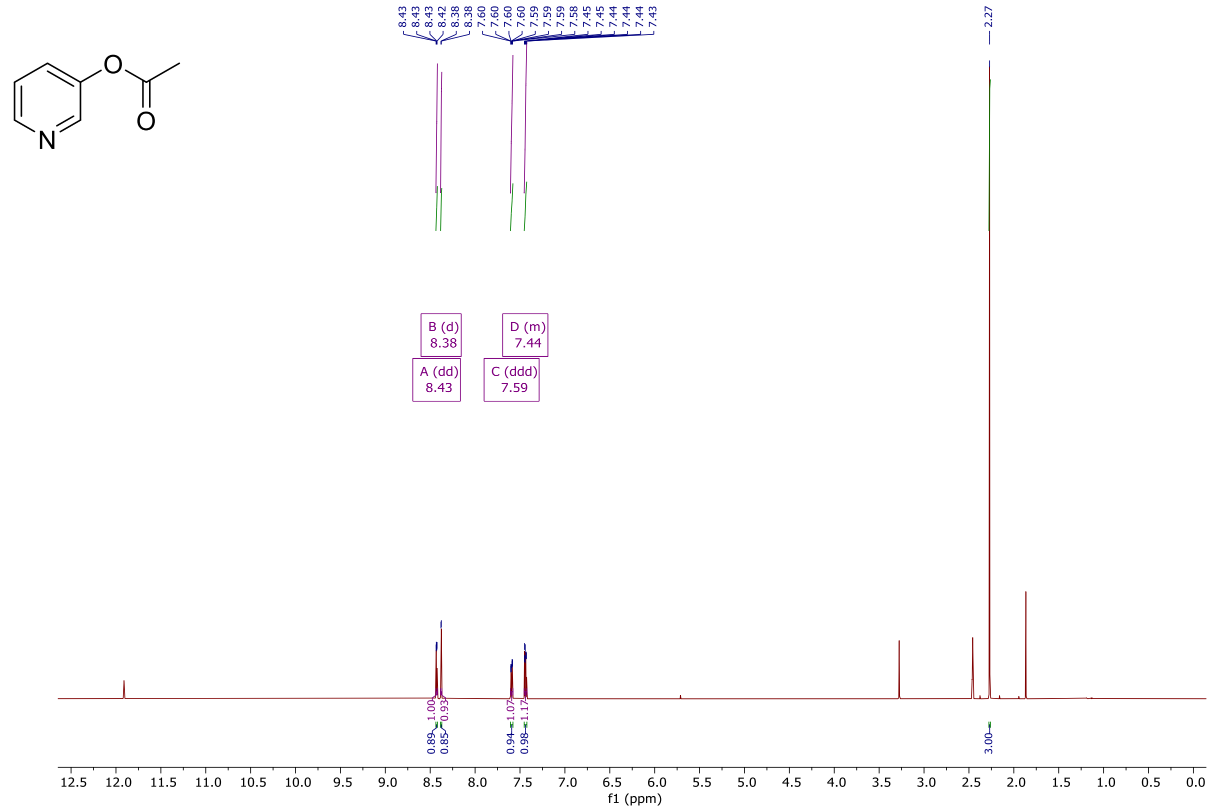

$^{1}$H NMR spectrum of compound **3** (600 MHz, DMSO-$d_6$)

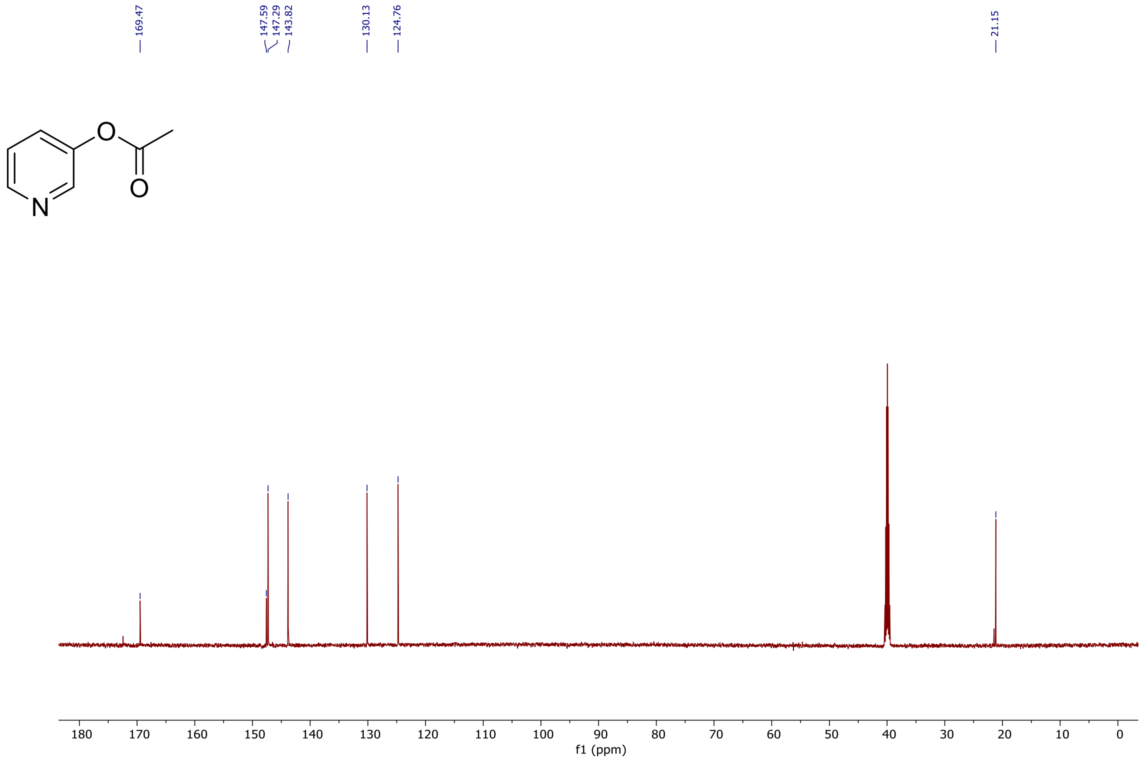

$^{13}$C NMR spectrum of compound **3** (150 MHz, DMSO-$d_6$)

tmcl_X240_35124.raw, C7H7NO2 [H]+

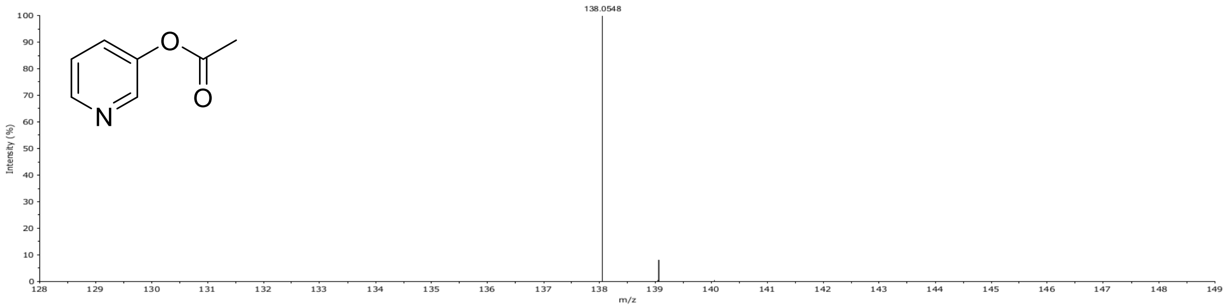

HRMS spectrum of compound **3**

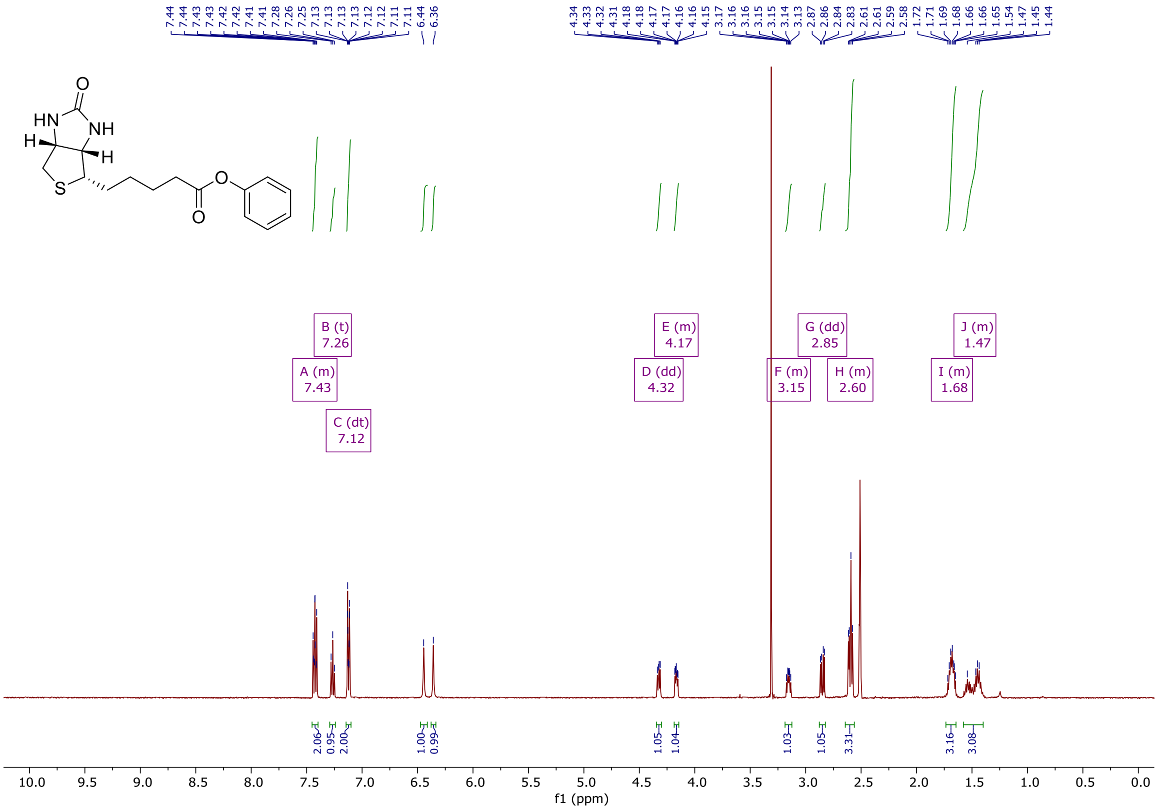

$^1$H NMR spectrum of compound **4** (500 MHz, DMSO-$d_6$)

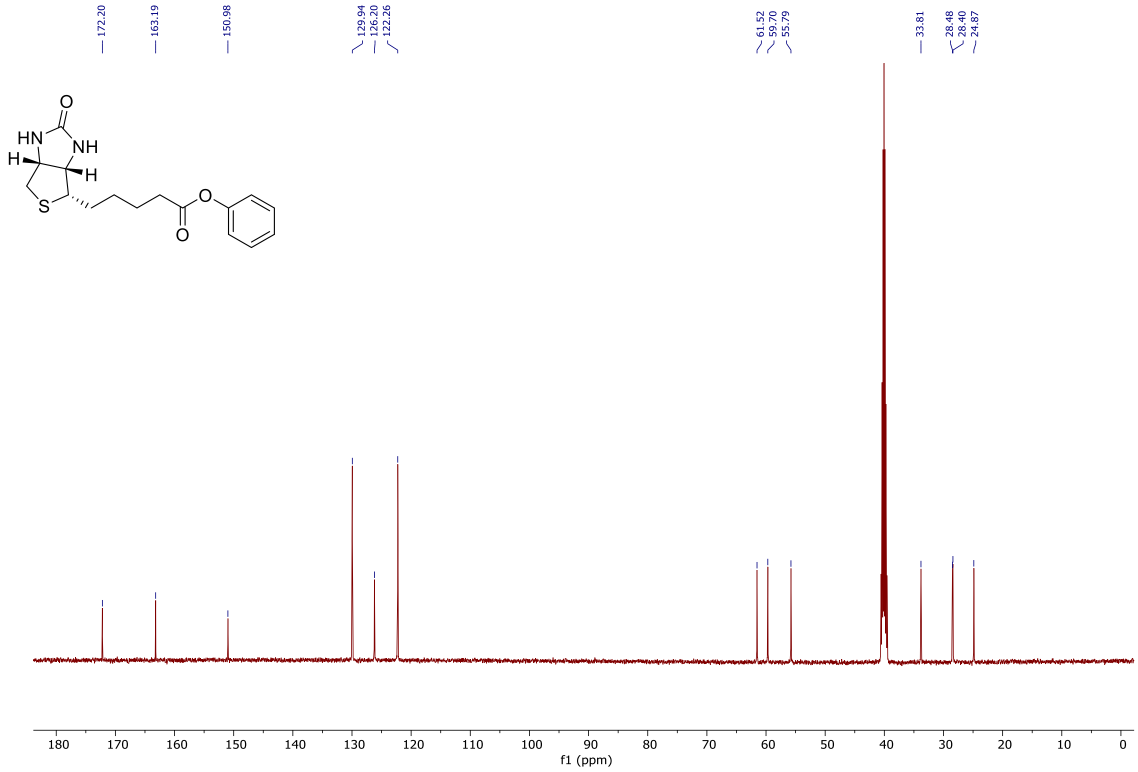

$^{13}$C NMR spectrum of compound **4** (125 MHz, DMSO-$d_6$)

tmcl_X240_35123.raw, C16H20N2O3S [H]+

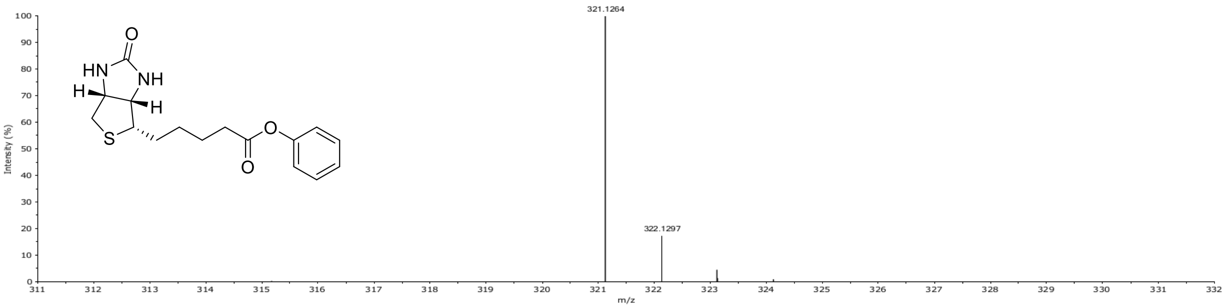

HRMS spectrum of compound **4**

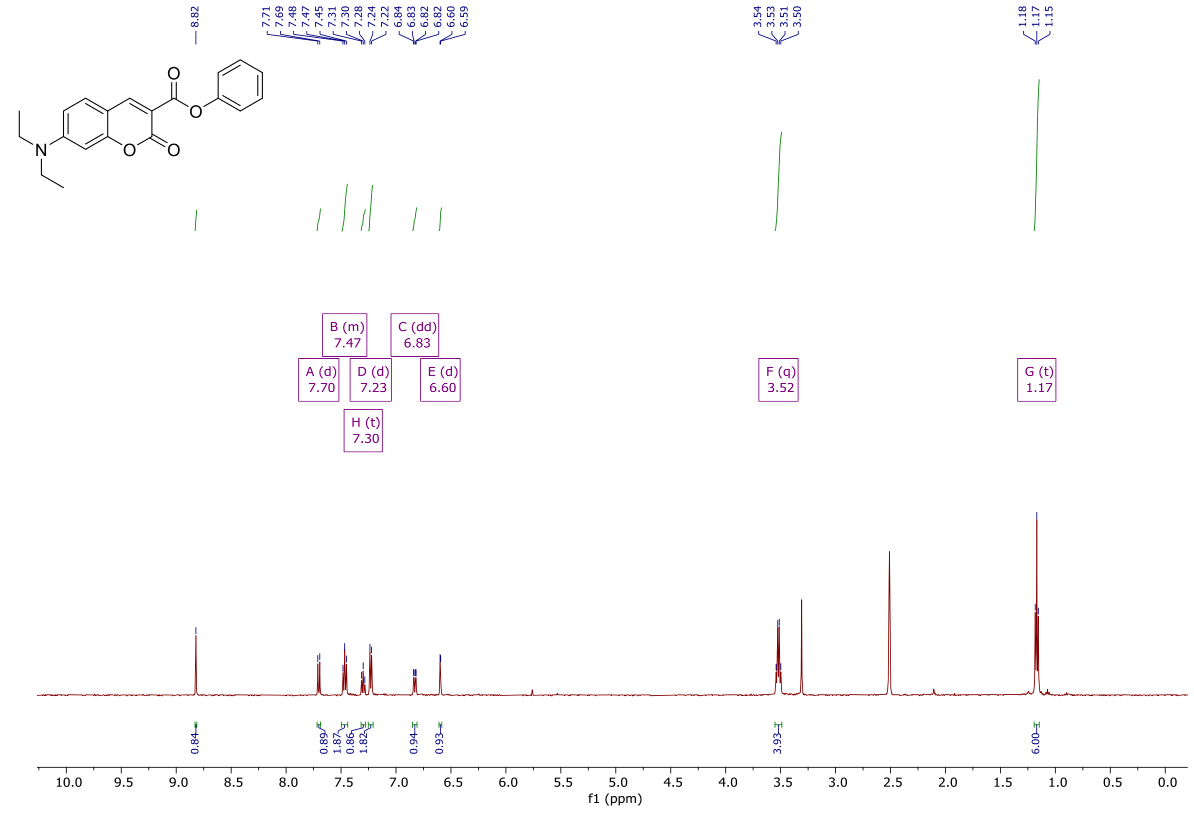

$^{1}H$ NMR spectrum of compound **5** (500 MHz, DMSO-$d_6$)

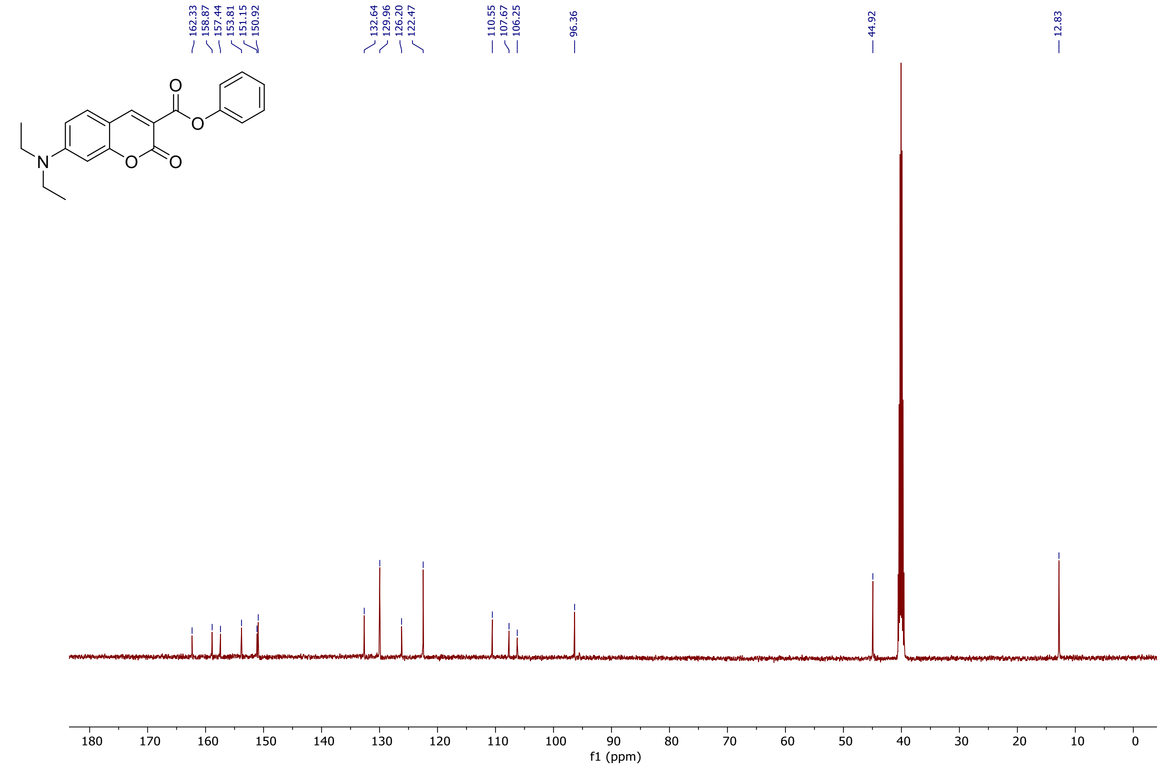

$^{13}C$ NMR spectrum of compound **5** (125 MHz, DMSO-$d_6$)

tmcl_X240_35609.raw, C20H19NO4 [H]+

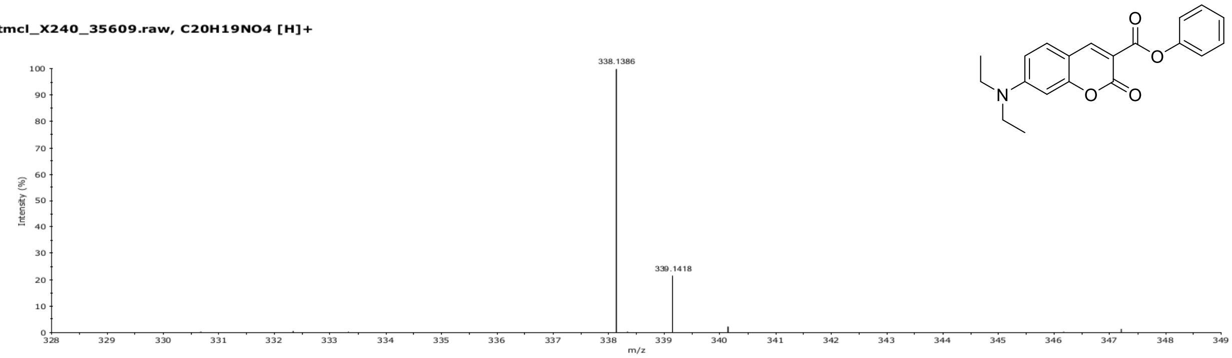

HRMS spectrum of compound **5**
